# A Novel Single Vector Intersectional AAV Strategy for Interrogating Cellular Diversity and Brain Function

**DOI:** 10.1101/2023.02.07.527312

**Authors:** Alex C. Hughes, Brittany G. Pollard, Beisi Xu, Jesse W. Gammons, Phillip Chapman, Jay B. Bikoff, Lindsay A. Schwarz

## Abstract

As the discovery of cellular diversity in the brain accelerates, so does the need for functional tools that target cells based on multiple features, such as gene expression and projection target. By selectively driving recombinase expression in a feature-specific manner, one can utilize intersectional strategies to conditionally promote payload expression only where multiple features overlap. We developed Conditional Viral Expression by Ribozyme Guided Degradation (ConVERGD), a single-construct intersectional targeting strategy that combines a self-cleaving ribozyme with traditional FLEx switches. ConVERGD offers benefits over existing platforms, such as expanded intersectionality, the ability to accommodate larger and more complex payloads, and a vector design that is easily modified to better facilitate rapid toolkit expansion. To demonstrate its utility for interrogating neural circuitry, we employed ConVERGD to target an unexplored subpopulation of norepinephrine (NE)-producing neurons within the rodent locus coeruleus (LC) identified via single-cell transcriptomic profiling to co-express the stress-related endogenous opioid gene prodynorphin (*Pdyn*). These studies showcase ConVERGD as a versatile tool for targeting diverse cell types and reveal *Pdyn*-expressing NE^+^ LC neurons as a small neuronal subpopulation capable of driving anxiogenic behavioral responses in rodents.

## Introduction

The ability to study the organization and function of neural circuits relies critically on knowledge of their underlying cellular diversity. Neurons have traditionally been characterized by their use of specific neurotransmitters or expression of specific genes. However, advances in sequencing technologies have supported a rapid expansion in cellular taxonomy that highlights the importance of considering more complex molecular diversity when defining neuronal cell types^1–4^. To target specific cell types in the rodent, researchers often rely on transgenic lines where recombinase-encoding genes are inserted into the genome near endogenous promoters or loci that direct their expression. Pairing these lines with transgenic or viral recombination-dependent reporter tools allows one to limit payload expression to defined cellular populations. Yet, as our understanding of cellular diversity in the brain expands, so does the need for more sophisticated tools that can selectively target cells based on multiple features. To address this need, several transgenic^5–7^ and viral-based^8–13^ intersectional strategies have been developed, each with unique strengths and weaknesses. Compared to viral strategies, transgenic approaches can better accommodate reporter transgenes that utilize large stop cassettes and may also be preferable for driving intersectional expression during development or in cell populations that are difficult to access^5–7^. On the other hand, transgenic reporters typically lack regional and temporal specificity, thus requiring careful consideration of off-target expression. These tools are also expensive and time-consuming to generate. Adeno-associated virus (AAV)-based reporter tools are an alternative approach that offer regional and temporal specificity and are relatively quick and inexpensive to generate. Because of these benefits, several elegant intersectional AAV tools with varying levels of *in vivo* functionality have recently emerged^8–13^. While useful, these tools also present a variety of challenges for the end user that may discourage their widespread adoption, including complex design parameters, use of large stop-cassettes that impede viral packaging capacity limits, dependence on multiple vectors for payload expression, or reliance on tetracycline-based expression systems which have reported issues with toxicity or leak expression^14–17^.

To address these limitations, we have developed a novel, single AAV vector-based intersectional targeting approach called Conditional Viral Expression by Ribozyme Guided Degradation (ConVERGD). ConVERGD is unique from other conditional expression strategies in that it relies on hammerhead ribozyme (HHR)-mediated mRNA inactivation^18,19^. Flanking the ribozyme with recombinase sites enables recombination to stabilize transcribed mRNA in a conditional manner. The design of ConVERGD introduces several improvements for the end user. Chiefly, the simplicity of ConVERGD means that transgenes do not need to be split into multiple components to achieve intersectionality. In addition, the small size of the ribozyme enables ConVERGD to be paired with larger promoters and transgenes while still remaining within AAV packaging capacity limits. This space-saving feature also provides efficient payload silencing within a single vector, thus eliminating the need to inject multiple viral vectors into targeted tissues.

To test the functionality of ConVERGD *in vivo*, we focused on a population of norepinephrine (NE, also called noradrenaline)-producing neurons in the mouse brainstem known as the locus coeruleus (LC). Activation of LC neurons promotes a wide range of arousal-related behavioral states ranging from enhanced memory formation and attentiveness to stress-related responses and anxiety^20^. Yet, how these diverse behavioral states are promoted by LC neurons remains largely unexplored. Using single-cell RNA sequencing, we identified a distinct subset of LC neurons that co-express Prodynorphin (Pdyn), which encodes an endogenous opioid implicated in the promotion of aversive emotional states. To determine how Pdyn^+^/Dbh^+^ LC neurons contribute to the array of behaviors broadly associated with LC activity, we developed an arsenal of ConVERGD-based viral tools that allowed us to identify the anatomic connectivity and behavioral influence of this unstudied subpopulation of neurons for the first time. Additionally, these studies showcase ConVERGD as a novel and easily amenable strategy for selectively targeting cell types defined by multiple features *in vivo*.

## Results

### Developing ConVERGD

Intersectional AAV-based tools often rely on complex design parameters, bulky silencing cassettes, and the use of multiple viruses to achieve specific targeting^8–13^ which limits their ability to be easily modified or accommodate new and more complex payloads. To address this, we developed a novel construct with a unique combination of genetic repression methods that leaves the transgene coding region intact and saves space in the viral packaging capacity. ConVERGD combines a small, loxP-flanked, self-cleaving type III hammerhead ribozyme variant (T3H48)^19^ with a traditional Flp-dependent FLEx switch^21^ (FRT/FRT5) that inverts the transgene coding region (Fig. 1a). In the presence of Flp, the transgene is oriented to be in line with the promoter, thereby allowing transcription to occur. However, the mRNA is immediately destabilized by ribozyme-mediated cleavage, preventing translation and, therefore, protein production (Fig. 1b). Only when the ribozyme is excised via Cre-mediated recombination can the transcribed mRNA remain stable and be translated to the final protein product. We tested this strategy by transfecting mouse Neuro2a (N2a) cells with a ConVERGD construct expressing eGFP (pAAV-hSyn-ConVERGD-eGFP-W3SL) alone or with recombinase expression plasmids containing either Cre, Flp, or both Cre and Flp (pDIRE). Under these conditions, eGFP expression occurred only in cells co-transfected with ConVERGD-eGFP and pDIRE (Fig. 1c). These results were validated quantitatively via FACS analysis (Fig. 1d), confirming that ConVERGD-based plasmids yield intersectional expression only when in the presence of both Cre and Flp recombinases *in vitro*.

**Fig. 1.**
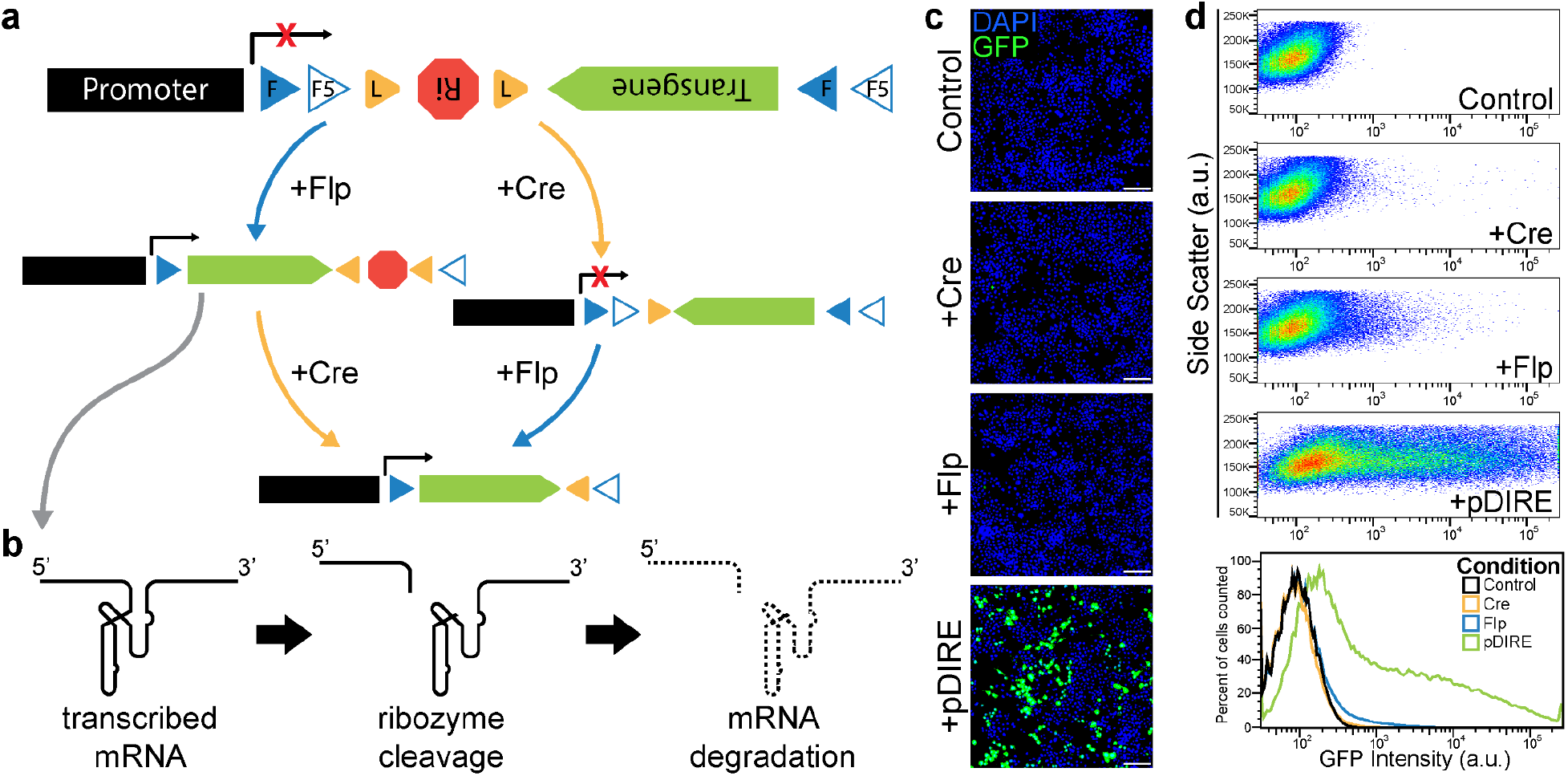
ConVERGD-based expression is dependent upon the presence of both Cre and Flp recombinases. **a**, Schematic of ConVERGD (Conditional Viral Expression by Ribozyme Guided Degradation). In the absence of recombinase or in the presence of Cre recombinase (yellow arrow), the transgene is not transcribed due to the inversion of the transgene coding region. In the presence of Flp recombinase (blue arrow), the transgene is transcribed but the mRNA is destabilized by ribozyme (Ri)-mediated cleavage and ultimately degraded (grey arrow). **b**, Schematic of ribozyme-mediated cleavage and degradation of transcribed mRNA. **c**, Representative images show hSyn-ConVERGD-eGFP (green) expression in DAPI (blue) counterstained N2a cell transfections. **d**, hSyn-ConVERGD-eGFP N2a cell transfections quantified by FACS analysis. Cell counts are normalized to the mode fluorescence value within each condition. Images in **c** are representative of results observed in 20 separate transfection experiments. FACS data in **d** represents pooled data collected across 6 separate transfection experiments. Scale bars are 100μm. F - FRT, F5 - FRT5, L - loxP, Ri - Ribozyme.

Our initial version of ConVERGD utilized the relatively small hSyn promoter and the optimized posttranscriptional regulatory element W3SL^22^ in order to conserve space for larger payloads. Still, the simple design and small size of the floxed ribozyme in ConVERGD facilitates its inclusion with other elements while still leaving sufficient space for large payloads. To test how ConVERGD performs when paired with various promoters (hSyn, CAG, EF1a, and nEF) and posttranscriptional regulatory elements (WPRE and W3SL), we designed additional ConVERGD vectors and tested their performance in N2a cells as described above. Two control conditions were included to directly compare ConVERGD with a single-recombinase vector (the Flp-dependent pAAV-hSyn-FLEx(FRT)eGFP-W3SL) and the original intersectional vector (the Cre- and Flp-dependent pAAV-hSyn-INTRSECT-Con/Fon-EYFP-WPRE^8^) (Extended Data Fig. 1). Cells co-transfected with ConVERGD-eGFP and pDIRE showed similar eGFP expression to that of cells co-transfected with FLEx(FRT)-eGFP and pDIRE. ConVERGD-eGFP also showed similar specificity compared to INTRSECT-Con/Fon-EYFP, indicating that ConVERGD’s *in vitro* performance is on par with current field-standard single-recombinase and intersectional constructs. Pairing ConVERGD with different posttranscriptional regulatory elements (WPRE or W3SL) did not influence expression (Extended Data Fig. 1). However, as expected, we observed an increase in fold induction in cells transfected with ConVERGD variants containing the stronger CAG, Ef1a, and nEF promoters (Extended Data Fig. 1). In these conditions, we also observed increased expression, albeit to a much lesser extent, in the Flp-alone conditions, suggesting that promoter strength could influence the ability of the ribozyme to sufficiently cleave all transcribed mRNA. Based on these observations, we focused subsequent toolkit expansion and *in vivo* experiments on ConVERGD constructs containing the hSyn promoter and W3SL post-translational regulatory element due to the small size and specific expression patterns of ConVERGD constructs containing these elements.

### Expanding ConVERGD Boolean logics and transgene expression

In addition to AND (CreANDFlp) Boolean logic, NOT logic (CreNOTFlp; FlpNOTCre) is useful in situations requiring access to intermingled cell populations that share the expression of certain genes but not others. We achieved ConVERGD NOT logic constructs by utilizing the inherent directionality of the ribozyme. We placed the transgene coding sequence and the ribozyme in opposite orientations and within FLEx switches so that the presence of one recombinase inverts both the transgene into the proper orientation for transcription and places the ribozyme in an inverted orientation so that it is not transcribed. In the presence of a second recombinase, the ribozyme is switched into a transcribable orientation, leading to mRNA destabilization and thus, preventing translation of the final protein product (Fig. 2a, c). ConVERGD-ConFoff (CreNOTFlp) (Fig. 2a-b) and ConVERGD-CoffFon (FlpNOTCre) (Fig. 2c-d) constructs both showed their highest expression levels in conditions where a single recombinase was present, with expression decreasing in the presence of a second recombinase, though not to control levels (Fig. 2b, d). We predict that the residual transgene expression observed in the presence of both recombinases is not unique to ConVERGD, but rather is caused by the imperfect recombination efficiency of Cre and Flp^23^, a phenomenon that was also reported for INTRSECT constructs, particularly in the CreNOTFlp orientation^8,9^.

**Fig. 2.**
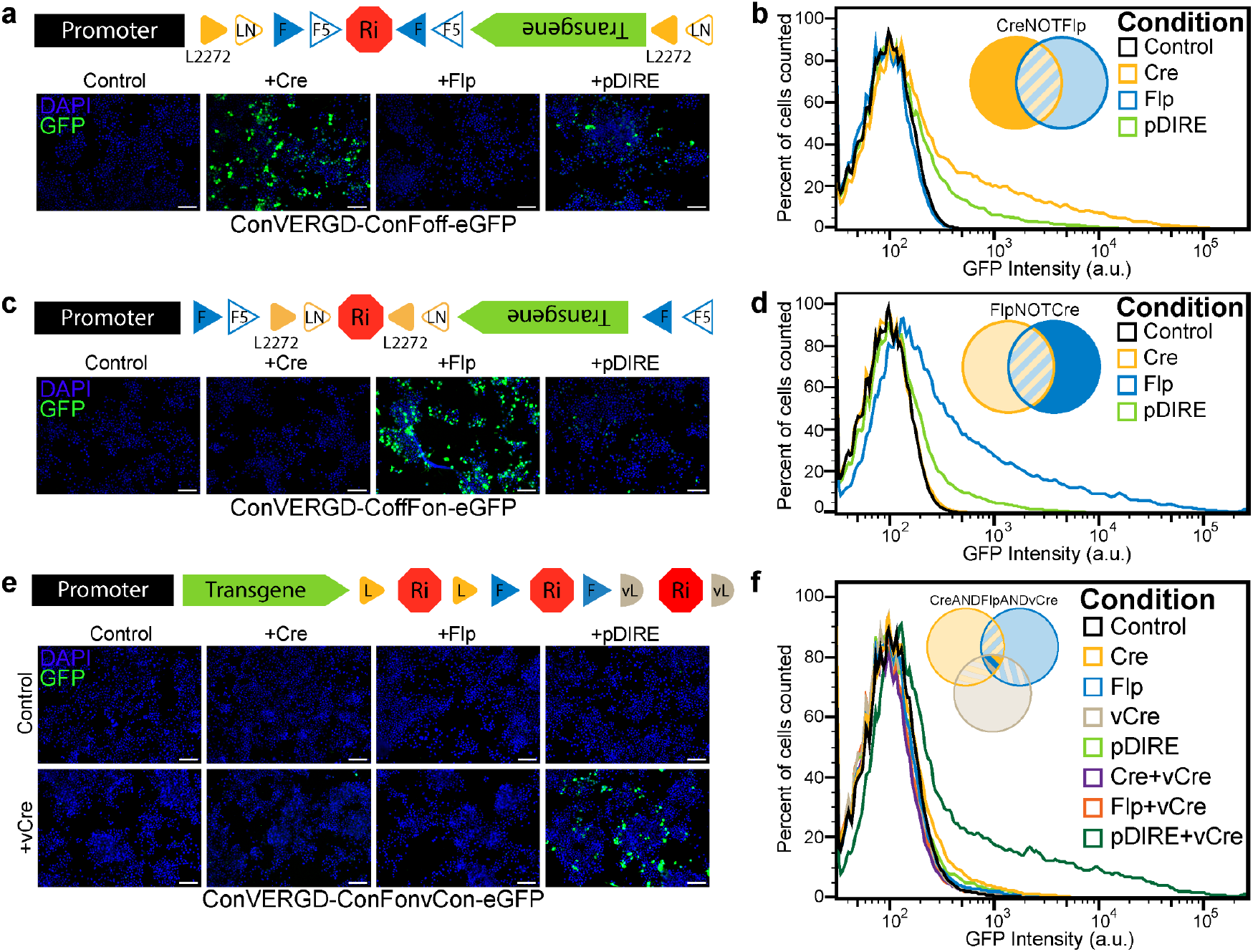
ConVERGD can be expanded to additional Boolean expression logics. **a**, ConVERGD-based ConFoff (CreNOTFlp) expression. By placing the self-cleaving ribozyme within a FLEx(FRT) switch that is antisense to the transgene coding region, ConVERGD constructs can be turned on by Cre and turned off by Flp. Representative images show hSyn-ConVERGD-ConFoff-eGFP (green) expression in DAPI (blue) counterstained N2a cell transfections. **b**, hSyn-ConVERGD-ConFoff-eGFP N2a cell transfections quantified by FACS analysis. Cell counts are normalized to the mode fluorescence value within each condition. **c**, ConVERGD-based CoffFon (FlpNOTCre) expression. By placing the self-cleaving ribozyme within a FLEx(loxP) switch that is antisense to the transgene coding region, ConVERGD constructs can be turned on by Flp and turned off by Cre. Representative images show hSyn-ConVERGD-CoffFon-eGFP (green) expression in DAPI (blue) counterstained N2a cell transfections. **d**, hSyn-ConVERGD-CoffFon-eGFP N2a cell transfections quantified as in **b. e**, ConVERGD-based triple-AND ConFonvCon (CreANDFlpANDvCRE) expression. Three uniquely flanked ribozymes were placed after the transgene coding region to achieve triple-intersectional expression that is dependent on the presence of Cre, Flp, and vCre. Representative images show hSyn-ConVERGD-ConFonvCon-eGFP (green) expression in DAPI (blue) counterstained N2a cells. **f**, hSyn-ConVERGD-ConFonvCon-eGFP N2a cell transfections quantified as in **b**. Images in **a, c**, and **e** are representative of results observed across 4 separate transfection experiments per construct. FACS results in **b, d**, and **f** represent pooled data from 4, 5, and 4 separate transfection experiments, respectively. Scale bars are 100μm. F - FRT, F5 - FRT5, L - loxP, vL - vloxP, Ri - Ribozyme.

Currently, intersectional strategies largely exist as dual-feature tools, a property that may be limiting as biological studies across scientific fields move towards characterizing more complex cell types. The INTRSECT toolkit recently developed a triple intersectional construct, called Triplesect, that allows payload expression in the presence of three orthogonal recombinases^9^. However, like all INTRSECT-based vectors, expression specificity is achieved via a complex genomic engineering pipeline that is unique to each payload and challenging to implement. We reasoned that the small size and simplicity of ConVERGD could provide an amenable framework for expanding a multi-feature intersectional toolkit beyond current strategies. To this end, we designed triple, quadruple, and quintuple intersectional ConVERGD variants. ConVERGD-ConFonvCon (CreANDFlpANDvCre), -ConFonvConNon (CreANDFlpANDvCreANDNigri), and -ConFonvConNonVon (CreANDFlpANDvCreANDNigriANDVika) constructs were created by placing three, four, or five ribozymes, respectively, each flanked with unique recombinase sites, after the transgene, so that activity from all recombinases is needed to excise all ribozymes and stabilize the mRNA. *In vitro* experiments confirmed the functionality of these vectors (Fig. 2e-f; Fig. 3a-b). While expanding the degree of intersectionality, these ConVERGD constructs also require less genetic space than Triplesect (Extended Data Table 1), a design feature that allows these tools to be paired with larger transgenes and promoters. Together, our data show that the inherent directionality and small size of the ribozyme can be utilized to create ConVERGD constructs with multiple expression logics (dual-feature AND, dual-feature NOT, triple-, quadruple-, and quintuple-feature AND) that expand intersectionality beyond what is possible with current tools.

**Fig. 3.**
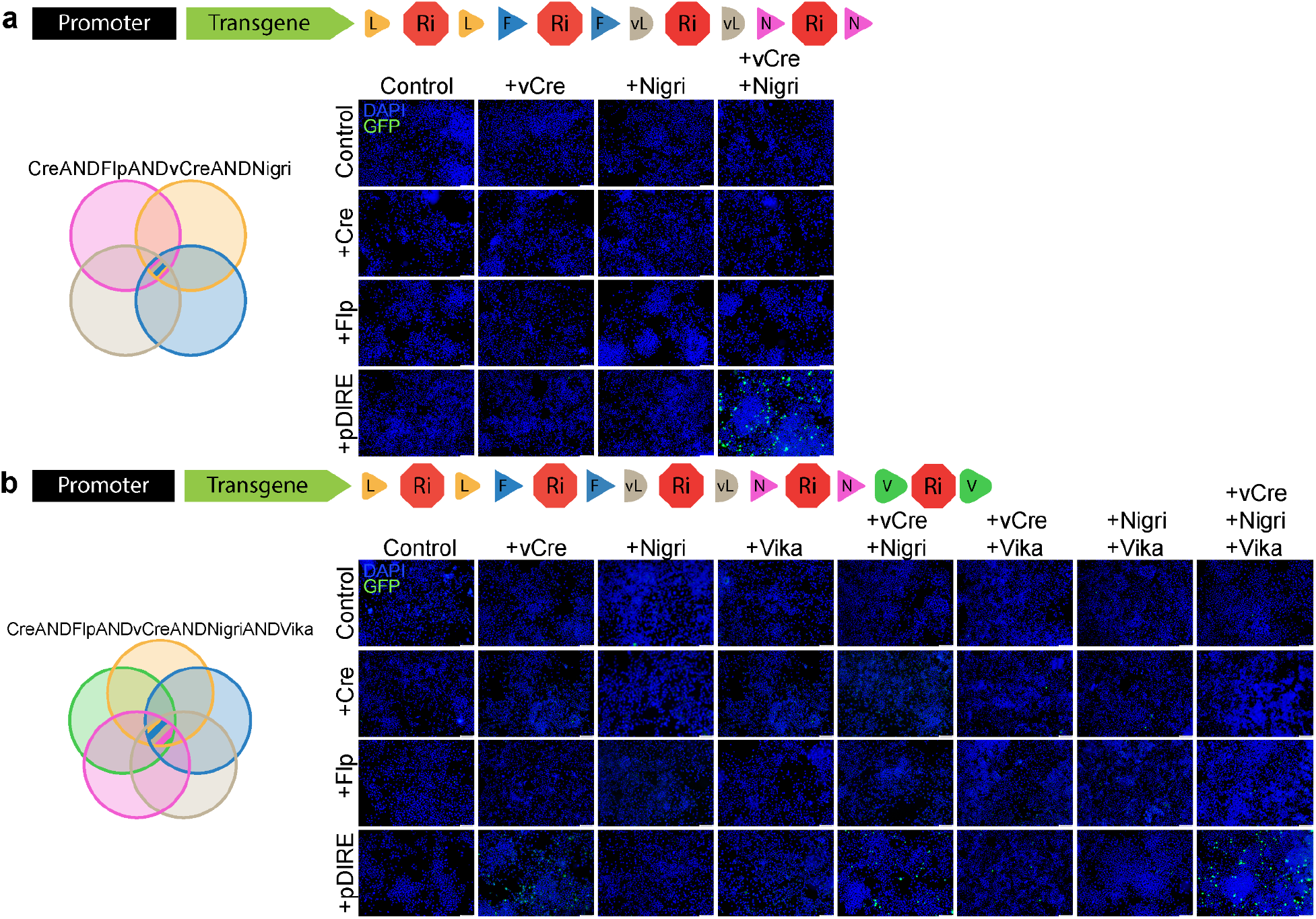
ConVERGD can be expanded to quadruple and quintuple intersectionality. **a**, ConVERGD-based quadruple-AND ConFonvConNon (CreANDFlpANDvCreANDNigri) expression. Four uniquely flanked ribozymes were placed after the transgene coding region to achieve quadruple-intersectional expression that is dependent on the presence of Cre, Flp, vCre, and Nigri.**b**, ConVERGD-based quintuple-AND ConFonvConNonVon (CreANDFlpANDvCreANDNigriANDVika) expression. Five uniquely flanked ribozymes were placed after the transgene coding region to achieve quintuple-intersectional expression that is dependent on the presence of Cre, Flp, vCre, Nigri, and Vika. All images show transfected N2a cells counterstained with DAPI (blue) and are representative of results observed across at least two separate transfections. Scale bars are 100μm. F - FRT, F5 - FRT5, L - loxP, vL - vloxP, N - Nox, V - Vox, Ri - Ribozyme.

Finally, the CreANDFlp ConVERGD toolkit was expanded to include several constructs relevant for *in vivo* study of neural circuits, such as chemogenetic activators and inhibitors (hM3Dq and hM4Di), optogenetic activators and inhibitors (ChR2, ArchT, and stGtACR), components for trans-synaptic rabies tracing (TVA receptor and N2c glycoprotein), calcium imaging tools (GCaMP8m), and a dual-labeling construct to visualize axonal projections and pre-synaptic sites (synaptophysin-GreenLantern-T2A-GAP43-mScarlet) (Extended Data Fig. 2). While these constructs represent only a small fraction of tools that are commonly used in neuroscience research, they highlight the amenability of ConVERGD to promote intersectional expression of diverse and complex transgenes with minimal design considerations.

### *In vivo* validation of ConVERGD viral vectors

ConVERGD vectors were next tested *in vivo* through targeted viral injections into the LC of transgenic mice where the NE-synthesizing enzyme dopamine-β-hydroxylase (*Dbh*) drives expression of either Flp or Cre recombinase. *Dbh*^*Flp*^ mice received an injection of AAV-hSyn-ConVERGD-eGFP into one LC nucleus and a mixture of ConVERGD-eGFP and AAV-EF1a-mCherry-IRES-Cre into the other LC nucleus (Fig. 4a). Injected *Dbh*^*Flp*^ mice showed eGFP expression only in LC neurons that also received Cre-expressing AAV (Fig. 4a). Similar results were achieved when ConVERGD-eGFP and AAV-EF1a-mCherry-IRES-Flp were co-injected into the LC of *Dbh*^*Cre*^ mice; eGFP expression occurred only in the LC nucleus co-injected with ConVERGD-eGFP and Flp (Fig. 4b). These results indicate that ConVERGD vectors support intersectional payload expression *in vivo* selectively in cells where Cre and Flp recombinase are present.

**Fig. 4.**
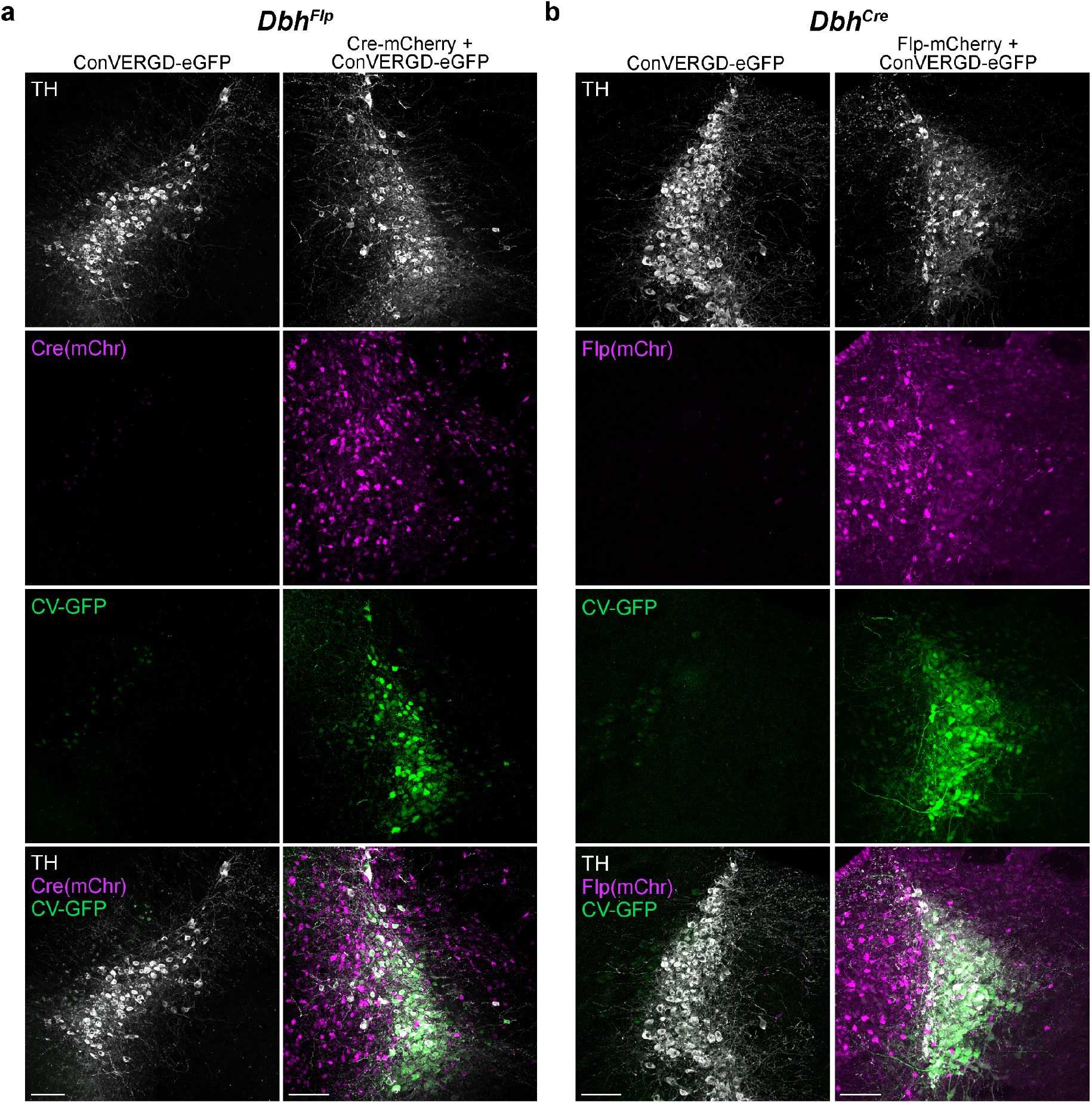
ConVERGD shows specific expression *in vivo* only in the presence of Cre and Flp recombinases. **a**, Representative images of the LC (TH, white) of a *Dbh*^*Flp*^ mouse injected with ConVERGD-eGFP (green) into one LC nucleus (left column) and ConVERGD-eGFP + Cre-mCherry (Cre(mChr), magenta) into the other LC nucleus (right column). **b**, Representative images of the LC (TH, white) of a *Dbh*^*Cre*^ mouse injected with ConVERGD-eGFP (green) into one LC nucleus (left column) and ConVERGD-eGFP + Flp-mCherry (Flp(mChr), magenta) into the other LC nucleus (right column). Images in **a** and **b** are representative of results observed across 4 and 2 animals, respectively. Working titers of AAV-ConVERGD-eGFP were the same across all injections. Scale bars are 100μm. TH - tyrosine hydroxylase; mChr - mCherry; LC - locus coeruleus; *Dbh* - dopamine-β-hydroxylase.

### Single-cell RNA sequencing of LC neurons reveals heterogeneous co-expression of chemical messengers and signaling receptors

To assess ConVERGD’s utility for characterizing discrete cell types *in vivo*, we again turned to LC neurons, a neuromodulatory cell type whose molecular diversity is understudied. While the LC has traditionally been classified as a uniform area due to its high density of NE-producing neurons, expression of other neuromodulators with important roles in the brain, such as galanin and NPY, have also been reported in this region^24–28^. These findings suggest that LC neurons could alter their functional impact through the co-release of additional signaling molecules, though the extent to which this molecular heterogeneity exists, and how it relates to the organization and function of LC-related neuronal circuits, is not comprehensively understood. To address this, we first performed single-cell transcriptome profiling (sc-RNAseq) of adult mouse LC neurons. *Dbh*^*Cre*^ mice were crossed with the Cre-dependent transgenic reporter mouse *Ai14*^*29*^ to fluorescently label LC neurons for isolation. Cells were obtained using a manual approach where fluorescently labeled cells were removed directly from brain sections collected from adult male and female *Dbh*^*Cre*^;*Ai14* transgenic mice (Fig. 5a). Smart-seq2 was used to generate scRNA-seq libraries and sequencing was performed to a depth of >1 million uniquely mapped reads per cell (median 16.7 million, mapping rate 94.6%), resulting in a median of 9000 genes detected per cell. Contaminated samples (14 cells), samples with less than 1 million uniquely mapped reads (22 cells), and samples without *Dbh* expression (34 cells) were excluded from further analyses (Fig. 5b). This resulted in 201 LC neurons for which normalized gene expression was reviewed to identify enriched genes, both in a non-biased manner (Extended Data Fig. 3) and using specified Gene Ontology (GO) terms to identify transcripts associated with neuronal signaling (Fig. 5c). In addition to *Dbh*, samples had robust expression of tyrosine hydroxylase (*TH*, an enzyme in the NE-synthesizing pathway) and solute carrier family 6 member 2 (*Slc6a2*, a NE transporter), supporting their classification as NE^+^ neurons. We also observed strong expression of galanin across a majority of LC cells, consistent with other reports^25,28^ (Fig. 5c). scRNA-seq revealed co-expression of several other neurotransmitter-synthesizing and neuropeptide genes within *Dbh*-expressing samples, such as calcitonin-related polypeptide alpha (*Calca*) which encodes the calcitonin gene-related neuropeptide (CGRP), and gamma-aminobutyric acid receptor subunit theta (*Gabrq*), a mediator of inhibitory synaptic transmission. Smaller subsets of LC neurons expressed glutamate decarboxylase 2 (*Gad2*), an enzyme involved in the synthesis of the inhibitory neurotransmitter GABA, cocaine and amphetamine regulated transcript protein (*Cartpt*), a preprotein known to modulate feeding, reward, and stress response, and prodynorphin (*Pdyn*) and proenkephalin (*Penk*), genes encoding precursor peptides for the endogenous opioids dynorphin and enkephalin (Fig. 5c). Fluorescence in situ hybridization (FISH) was performed to explore the spatial location and validate co-expression of target genes within the *Dbh*^*+*^ LC cells. Consistent with our sequencing data, we observed that almost all *Dbh*-expressing LC neurons contained *Calca* and *Gabrq* transcripts, while more discrete subsets of LC cells co-expressed *Cartpt, Gad2, Pdyn*, and *Penk* (Fig. 5d). Collectively, these data show that LC neurons heterogeneously express transcripts encoding a variety of neurotransmitters and neuropeptides in addition to NE, potentially diversifying their functional impact in downstream brain areas.

**Fig 5.**
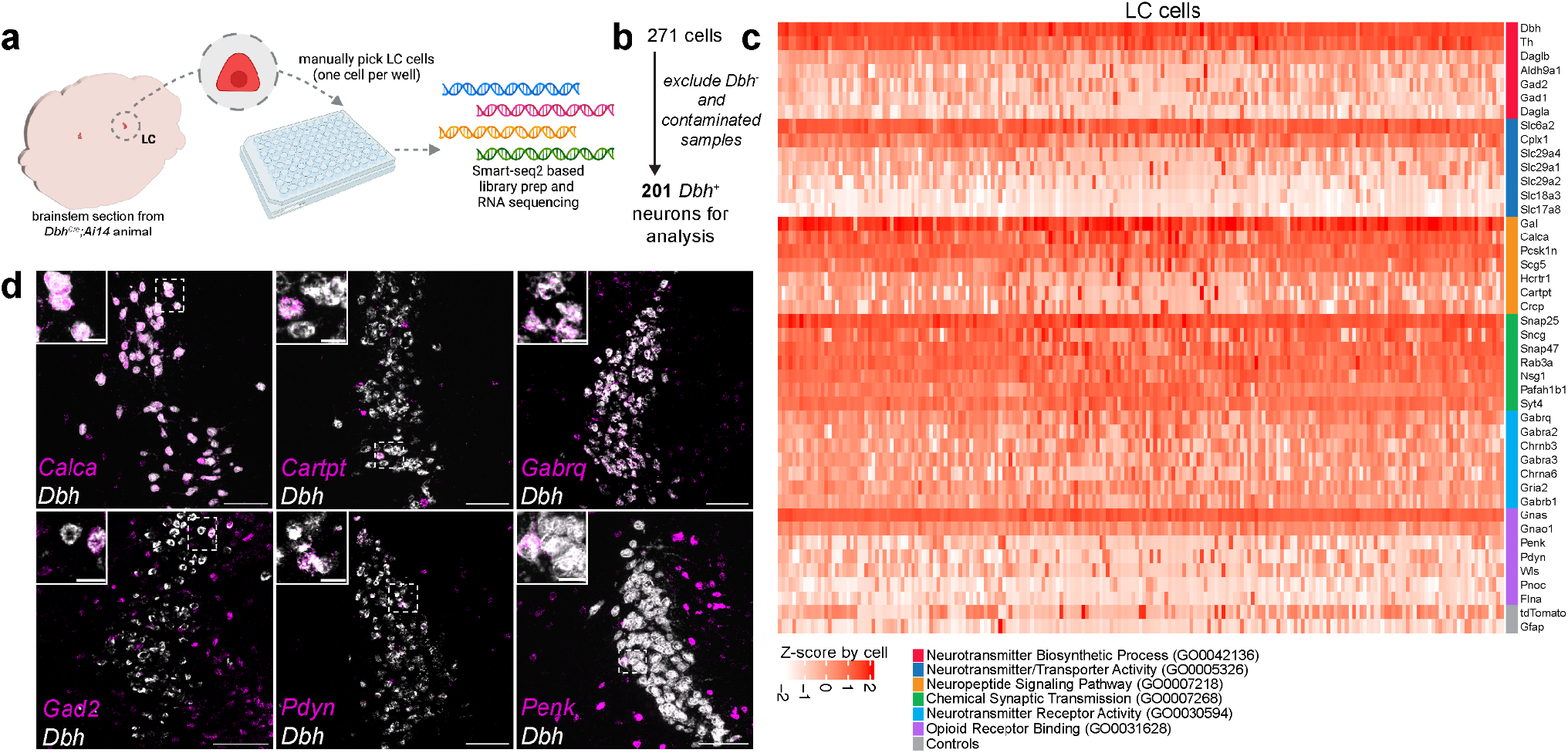
Subsets of norepinephrine LC neurons co-express genes for other neurotransmitters and neuropeptides. **a**, Workflow for molecular profiling of individual LC neurons. Fluorescently labeled LC neurons were manually isolated from adult mouse brain tissue for sequencing (n=14 mice). Full-length cDNA and sequencing libraries were generated for each cell using a Smart-seq2 based protocol. **b**, Of 271 cells, 201 were identified as norepinephrine (NE) LC neurons due to the presence of mRNA for the NE-synthesizing gene dopamine-β-hydroxylase (*Dbh*). **c**, Heatmap of scaled transcript abundance (transcripts per million, TPM) for LC neurons. The top expressing genes from GO term categories for neurotransmitters (red), transporters (dark blue), neuropeptides (orange), synaptic transmission (green), neurotransmitter receptors (light blue), opioid signaling (purple), or selected control genes (gray) are displayed. **d**, *In situ* hybridization of selected neuronal signaling molecules (magenta) within *Dbh*-expressing LC neurons (white). Transcripts for calcitonin-related polypeptide alpha (*Calca*), cocaine- and amphetamine-regulated transcript protein (*Cartpt*), gamma-aminobutyric acid type A receptor subunit theta (*Gabrq*), glutamate decarboxylase 2 (*Gad2*), prodynorphin (*Pdyn*), and proenkephalin (*Penk*) were expressed within LC cells. Scale bars are 100μm for larger images and 20μm for insets. LC - locus coeruleus

### Validation of ConVERGD to access prodynorphin-expressing LC neurons

Activation of the LC promotes diverse behavioral responses^20^, but how this nucleus, with its broad connectivity, maintains such a rich functional repertoire remains an open question in the field. One option is that subsets of LC neurons could form distinct neural circuits with specialized functions, an idea supported by recent work which showed that selective activation of certain LC projections promotes anxiogenic or anxiolytic-like behaviors^30–34^. Our sequencing data revealed a subpopulation of LC cells that co-express both *Dbh* and *Pdyn*, a neuropeptide generally associated with aversive behavioral responses related to stress^35–42^. As activation of the LC is also critical for modulating stress-related responses^20,43^, we were interested to learn more about the anatomical connectivity and functional relevance of this LC subpopulation using intersectional strategies. We first confirmed that transgenic animals could be used to target Pdyn^+^ LC neurons by crossing *Pdyn*^*Cre*^ mice with a Cre-dependent nuclear eGFP reporter line (*Sun1-eGFP*). GFP^+^ cells were observed adjacent to and within the LC in a pattern consistent with *Pdyn* expression reported in the Allen Institute for Brain Science mouse brain *in situ* datasets^44^. A subset of GFP^+^ cells in the area of the LC colocalized with immunolabeling against TH, confirming their identity as NE^+^ neurons (Fig. 6a). The spatial proximity of Pdyn^+^/TH^+^ LC neurons with nearby Pdyn^+^/TH^-^ cells (Fig. 6a, inset) precludes the use of single recombinase-dependent tools to target these populations independently, again justifying the need for intersectional strategies for further characterization.

**Fig. 6.**
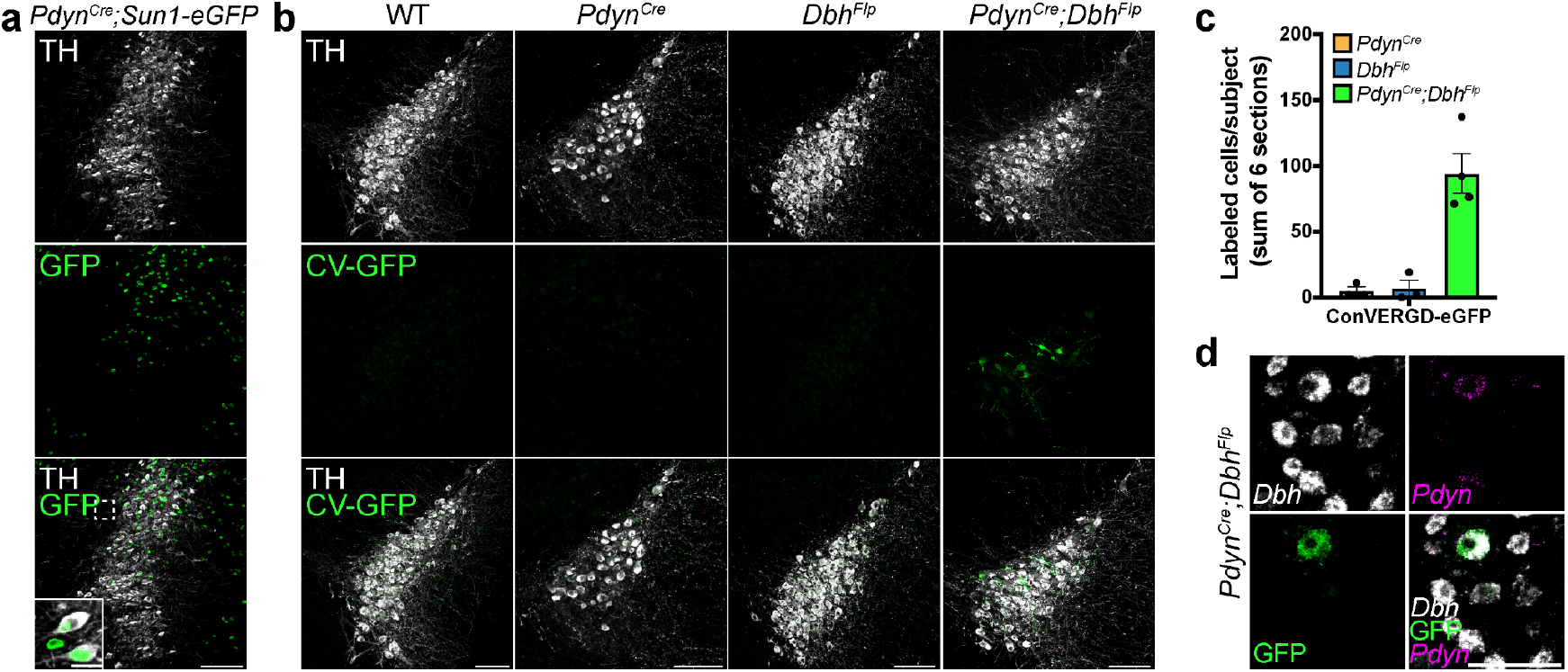
ConVERGD shows specific expression in *Pdyn*^*Cre*^*;Dbh*^*Flp*^ mice. **a**, Representative images of GFP (green) labeled Pdyn^+^ cells in and around the LC (TH, white) in the transgenic line *Pdyn*^*Cre*^*;Sun1-eGFP*. **b**, Representative images the LC (TH, white) of AAV-hSyn-ConVERGD-eGFP injected WT, *Pdyn*^*Cre*^, *Dbh*^*Flp*^, and *Pdyn*^*Cre*^*;Dbh*^*Flp*^ mice. GFP expression (green) was found only in a subset of LC cells in *Pdyn*^*Cre*^*;Dbh*^*Flp*^ mice. **c**, Quantification of ConVERGD-eGFP labeled cells in and around (∼200μm radius) the LC across different genotypes. Points represent the averaged cell counts across 6 50μm LC brain sections. Bars represent the mean of the data. Error bars are SEM. n = 3 (*Pdyn*^*Cre*^), 3 (*Dbh*^*Flp*^), and 4 (*Pdyn*^*Cre*^*;Dbh*^*Flp*^). **d**, Representative images of multiplexed labeling of ConVERGD-eGFP expression (green, immunostain) within a cell expressing *Dbh* (white, *in situ* hybridization) and *Pdyn* (magenta, *in situ* hybridization) within a ConVERGD-eGFP-injected *Pdyn*^*Cre*^*;Dbh*^*Flp*^ brain section. All sections from virally injected mice were immunostained against GFP. Images in **b** are representative of results observed across 2 (WT), 3 (*Pdyn*^*Cre*^), 3 (*Dbh*^*Flp*^), and 4 (*Pdyn*^*Cre*^*;Dbh*^*Flp*^) animals. Images in **d** are representative of results across two multiplexing experiments. Scale bars are 100μm for the larger images in **a-b**, and 20μm for the inset in **a**, and 50μm in **d**. TH - tyrosine hydroxylase; LC - locus coeruleus; WT - wild-type; *Pdyn* - prodynorphin; *Dbh* - dopamine-β-hydroxylase; CV - ConVERGD.

We next assessed if ConVERGD could selectively target Pdyn^+^ LC neurons by injecting AAV-hSyn-ConVERGD-eGFP into the LC of *Pdyn*^*Cre*^*;Dbh*^*Flp*^ mice or control animals (wild-type (WT), *Pdyn*^*Cre*^, and *Dbh*^*Flp*^ mice). While no GFP expression was observed in control mice, we observed a subset of labeled LC cells specifically in the *Pdyn*^*Cre*^*;Dbh*^*Flp*^ mice, indicating AAV payload expression is specific to Cre- and Flp-expressing cells (Fig. 6b-c). Multiplexed labeling of LC neurons expressing ConVERGD-eGFP revealed co-expression of *Pdyn* and *Dbh* mRNA in GFP-containing cells (Fig. 6d). ConVERGD’s performance *in vivo* was also compared directly to INTRSECT. AAV-hSyn-INTRSECT-Con/Fon-EYFP was injected into the LC of *Pdyn*^*Cre*^, *Dbh*^*Flp*^, and *Pdyn*^*Cre*^*;Dbh*^*Flp*^ mice at the same titer used for ConVERGD injections (∼7.87E11 gc/mL). Under these conditions, we observed a considerable number of INTRSECT labeled neurons in *Pdyn*^*Cre*^*;Dbh*^*Flp*^ samples. However, substantial labeling was also seen in *Pdyn*^*Cre*^ and *Dbh*^*Flp*^ samples, suggesting non-specific payload expression had occurred (Extended Data Fig. 4a-b). Collectively, these experiments establish ConVERGD as a viable intersectional approach for targeting discrete cell populations *in vivo* with improved selectivity over current field standard approaches.

### Pdyn^+^/Dbh^+^ LC neurons connect with brain regions implicated in stress response

We next sought to test the practical application of ConVERGD by investigating the connectivity of this molecularly distinct subpopulation of Pdyn^+^/Dbh^+^ LC neurons. Although several recent studies have highlighted the modular afferent and efferent patterns of LC neurons^20,43,45,46^, the input-output organization of molecularly distinct subpopulations of LC neurons remains understudied, partially due to the lack of sufficient tools to access these molecular subsets. Thus, we created ConVERGD-based helper AAVs containing a TVA receptor and the N2c glycoprotein and injected them into the LC of *Pdyn*^*Cre*^*;Dbh*^*Flp*^ mice to perform intersectional, CVS-N2c^ΔG^ rabies-based^47^, monosynaptic input tracing from Pdyn^+^/Dbh^+^ LC neurons (Fig. 7a). Within the LC, a subset of TH^+^ cells had overlapping ConVERGD-TVA(mCh) and RABV-CVS-N2c^ΔG^-H2B-eGFP labeling, indicative of ‘starter cells’ that support the trans-synaptic spread of the rabies virus to their presynaptic partners (Fig. 7b). Whole-brain quantification revealed distant inputs to Pdyn^+^/Dbh^+^ LC cells throughout the brain, with enriched labeling in certain regions (Fig. 7c). Inputs to Pdyn^+^/Dbh^+^ LC neurons were found in areas ranging from pontine regions such as the paragigantocellular reticular nucleus to frontal cortical regions such as the prelimbic cortical area. From this brain-wide labeling, sections containing at least 3% of all counted cells were selected for further interrogation to identify brain regions with enriched numbers of rabies-labeled inputs. From posterior to anterior, we found that the dorsal and lateral paragigantocellular reticular nucleus (PGRNd, PGRNl), midbrain reticular nucleus (MRN), periaqueductal gray (PAG), hypothalamus, central amygdala (CeA), bed nucleus of the stria terminalis (BNST), orbitofrontal cortex (OFC), and prelimbic cortical area (PLC) had the highest concentration of rabies-labeled presynaptic inputs (Fig. 6d). Although we did not fully dissect the distribution of input cells located in specific cortical areas, the strongest medial clusters fell within the prelimbic and motor cortical areas whereas the more lateral inputs came primarily from the OFC and the agranular insular area (AIA). Together, our data suggest that Pdyn^+^/Dbh^+^ LC cells receive inputs from regions spanning the anterior-posterior axis of the brain with the highest proportion of these afferents arising largely from stress-related brain areas such as the BNST, hypothalamus, and the CeA.

**Fig. 7.**
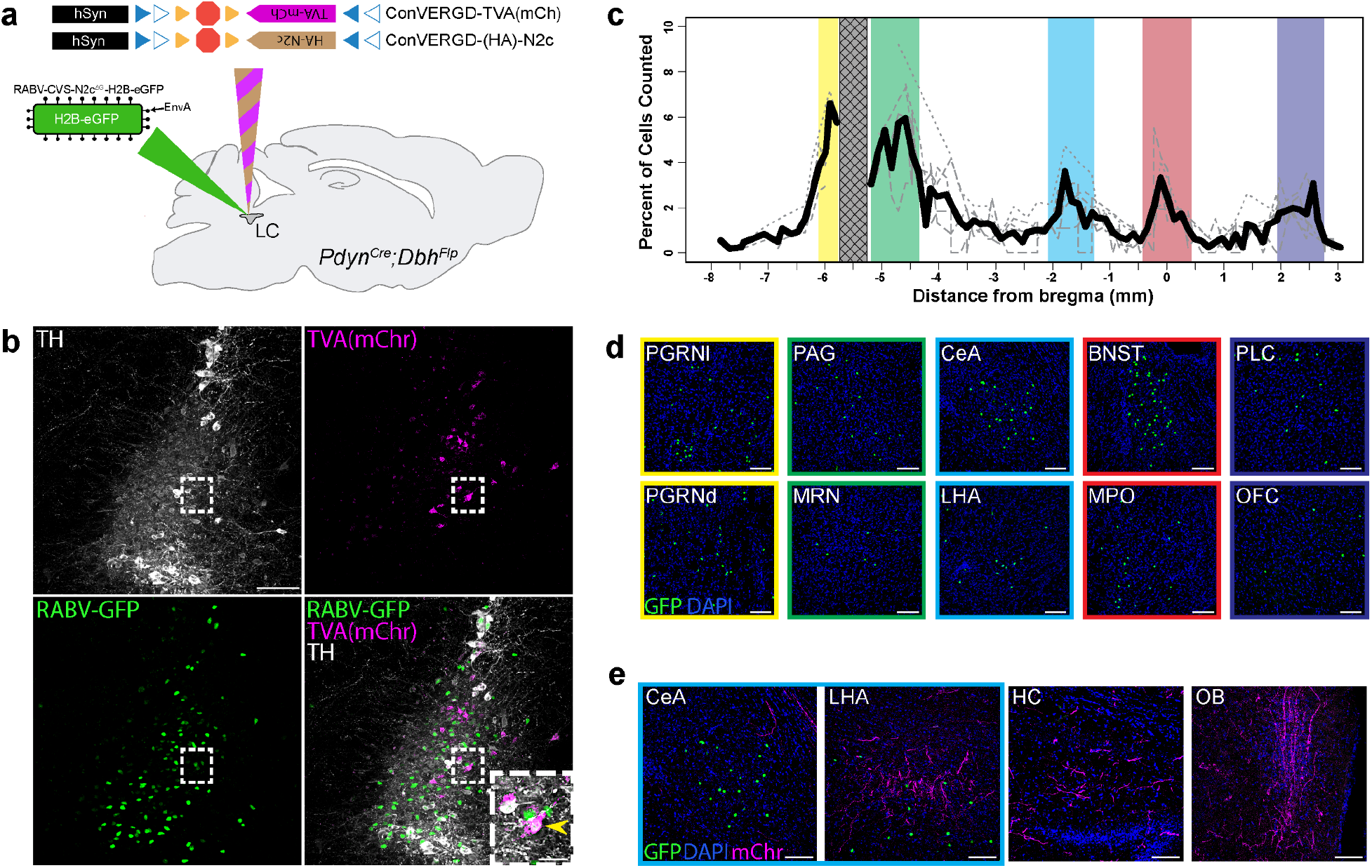
ConVERGD-based monosynaptic rabies tracing reveals inputs onto Pdyn^+^/Dbh^+^ LC cells. **a**, ConVERGD-based rabies helper AAVs expressing TVA receptor (magenta) and N2c glycoprotein (brown) were unilaterally co-injected into the LC of *Pdyn*^*Cre*^*;Dbh*^*Flp*^ mice three weeks prior to injection of RABV-CVS-N2c^ΔG^-H2B-eGFP (green). **b**, Representative images of TVA(mCh) expression (magenta) and RABV-GFP expression (green) within the LC (TH, white). TVA(mCh) is present only in a subset of LC cells. The inset image highlights a TH^+^ LC starter cell (yellow arrow) containing GFP expression from the rabies virus and mCherry expression from ConVERGD-TVA(mCh). **c**, Quantitative distribution of GFP^+^ cells across the brain. Highlighted regions represent areas containing at least 3% of total GFP^+^ cells counted. Colors correspond to the regions specified in **d**,. The grey cross-hatched region represents the area around the LC that was removed from quantification. Light gray traces represent the distribution of GFP^+^ cell labeling from individual brains. The bold line represents an average of all brains. n = 5 mice. **d**, Representative images of GFP^+^ cells in major input regions. Cell nuclei are counterstained with DAPI (blue). **e**, Representative images of axon projections from TVA(mCh)^+^ (magenta) LC cells. Nuclei are counterstained with DAPI (blue). PGRNl/d, paragigantocellular reticular nucleus lateral/dorsal region; PAG, periaqueductal gray; MRN, midbrain reticular nucleus; CeA, central amygdala; LHA, lateral hypothalamic area; BNST, bed nucleus of the stria terminalis; MPO, medial preoptic area of the hypothalamus; PLC, prelimbic cortical area; OFC, orbitofrontal cortex; HC, hippocampus; OB, olfactory bulb. Scale bars are 100μm in **b, d**, and **e**. Inset scale bar in **b**, is 25μm. TH - tyrosine hydroxylase; mChr - mCherry; LC - locus coeruleus; *Pdyn* - prodynorphin; *Dbh* - dopamine-β-hydroxylase.

The CVS-N2c^ΔG^ modified rabies virus was chosen for these studies because it has been reported to have improved tracing efficiency and reduced cellular toxicity over the B19 strain^47^. Indeed, we observed that the cellular integrity of the starter cells was well maintained, which, in turn, allowed us to trace the afferent axonal projections of Pdyn^+^/Dbh^+^ LC starter cells via their expression of TVA-mCherry to determine their projection targets. Of note, we observed regions, like the CeA and LHA, that received high levels of efferent projections but differed in their levels of afferent projections; the CeA had sparse innervation while the LHA was densely innervated by Pdyn^+^/Dbh^+^ LC cells (Fig. 7e). Strong projections were also found in areas that did not contain efferent projections, such as the hippocampus (HC) and the olfactory bulb (OB). Together, these data highlight nonuniform projection patterns of a molecularly distinct LC subpopulation, whereby Pdyn^+^/Dbh^+^ LC neurons preferentially innervate certain stress-associated brain structures but still broadly collateralize their axons to target many other brain regions.

### Activating Pdyn^+^/Dbh^+^ LC neurons drives an anxiogenic behavioral phenotype during elevated zero maze but not open field testing

Given that dynorphin signaling and LC activation have separately been implicated in promoting anxiogenic behaviors^20,27,30,34,48–51^ and that our connectivity data revealed enriched connections between Pdyn^+^/Dbh^+^ LC neurons and brain areas centrally involved in the stress response, we wondered how the selective activation of just Pdyn^+^/Dbh^+^ LC cells, which represent a small portion (>20%) of the LC, might influence behavior. To test this, we developed a ConVERGD-based excitatory DREADD construct to enable transient activation of Pdyn^+^/Dbh^+^ LC neurons during behavioral testing. Bilateral injection of AAV-hSyn-ConVERGD-(HA)hM3Dq into the LC of WT, *Pdyn*^*Cre*^, *Dbh*^*Flp*^, and *Pdyn*^*Cre*^*;Dbh*^*Flp*^ mice induced excitatory DREADD receptor expression in a subset of LC neurons only in *Pdyn*^*Cre*^*;Dbh*^*Flp*^ mice (Fig. 8a-b). We validated the functionality of hM3Dq expression by giving CNO intraperitoneally (IP) prior to perfusion and immunolabeling sections against the immediate early gene cFos to assess neuronal activation. *Pdyn*^*Cre*^*;Dbh*^*Flp*^ mice expressing ConVERGD-hM3D showed enhanced cFos expression in TH^+^ LC cells compared to *Pdyn*^*Cre*^ mice that had no hM3Dq expression (Fig. 8c). To test the behavioral effect of activating Pdyn^+^/Dbh^+^ LC neurons, we assessed performance during open field testing (OFT), an assay commonly used to quantify locomotion, exploration, and anxiety-like behavior in rodents^52^. All mice received bilateral injections of AAV-hSyn-ConVERGD-(HA)hM3Dq into the LC at least three weeks prior to behavioral testing and received CNO 30 minutes prior to placement in the open field, where distance and time spent in sheltered corner regions and the exposed center region were measured (Fig. 8d). Control mice included WT, *Pdyn*^*Cre*^, and *Dbh*^*Flp*^ animals after statistical analysis (ANOVA, see methods) revealed no behavioral differences between these genotypes. Mice in the control and experimental groups traveled similar total distances within the arena, with no difference in the time each group spent in the corners or center portion of the open field (Fig. 8e-g), indicating that selective activation of Pdyn^+^/Dbh^+^ LC neurons was not sufficient to drive behavioral alterations during OFT. We also activated Pdyn^+^/Dbh^+^ LC neurons while mice were placed in an elevated zero maze (EZM), an assay used to test anxiety in rodents^53^. As in OFT, mice were given CNO 30 minutes prior to being placed in the EMZ, where distance and time spent in closed and open regions were measured. Control and experimental groups traversed all areas of the EZM during their sessions (Fig. 8h) with no significant difference in the total distance traveled or distance traveled in the open regions (Fig. 8i-j). However, compared to control mice, hM3Dq-expressing *Pdyn*^*Cre*^*;Dbh*^*Flp*^ mice spent significantly less time in the open regions of the EZM compared to controls (Fig. 8k), suggesting an anxiogenic behavioral state portrayed by their reluctance to spend time in the unsheltered portions of the maze. Together, our behavioral tests show that activation of Pdyn^+^/Dbh^+^ LC cells, which represent a small percentage of cells within the LC, is sufficient to drive anxiety-like behaviors in EZM but not OFT. These findings differ from previous studies where activation of all LC neurons promoted anxiety-like behavior in both OFT and EZM^27,30,51,54^, and could suggest that activation of Pdyn^+^/Dbh^+^ LC neurons has a greater behavioral influence in acutely stressful situations, such as in the EZM.

**Fig. 8.**
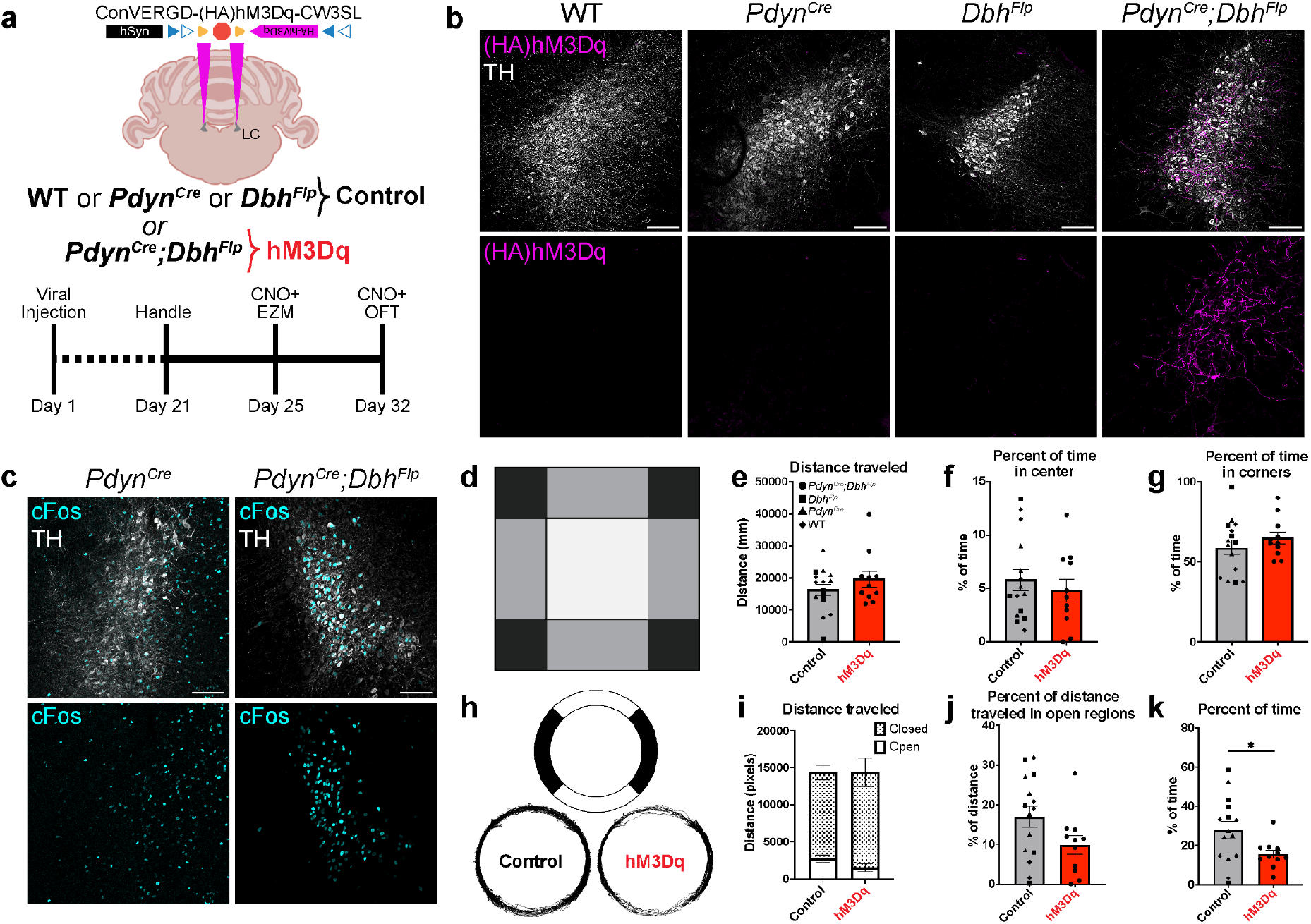
ConVERGD-based chemogenetic activation of Pdyn^+^/Dbh^+^ LC neurons promotes an anxiogenic phenotype in EZM but not OFT. **a**, Behavioral setup and timeline. AAV-hSyn-ConVERGD-(HA)hM3Dq was bilaterally injected into the LC of WT, *Pdyn*^*Cre*^, *Dbh*^*Flp*^, and *Pdyn*^*Cre*^*;Dbh*^*Flp*^ mice at least three weeks prior to behavioral tests. **b**, Representative images of hM3Dq expression (HA, magenta) observed in a subset of LC cells (TH, white) in *Pdyn*^*Cre*^*;Dbh*^*Flp*^ mice but not in control (WT, *Pdyn*^*Cre*^, *Dbh*^*Flp*^) mice. **c**, Representative images of cFos (cyan) expression following CNO injection in *Pdyn*^*Cre*^ and *Pdyn*^*Cre*^*;Dbh*^*Flp*^ mice injected with AAV-hSyn-ConVERGD-(HA)hM3Dq. **d**, Diagram of the open field chamber with quantified ROI center (white) and corner (black) divisions. Perimeter (grey) regions were not quantified. **e**, Total distance traveled during OFT. **f**, Percent of time spent in the center region of the open field chamber. **g**, Percent of time spent in the corners of the open field. **h**, Diagram of EZM (top) and combined traces of control and hM3Dq-expressing mice (bottom). **i**, Total distance (sum of closed (shaded) and open (white)) traveled in the EZM. **j**, Percent of distance traveled in the open region of the EZM. **k**, Percent of time spent in the open region of the EZM. hM3Dq-expressing mice spent significantly less time in the open regions compared to control mice. * p = 0.0314, two-tailed t-test. n = 15 Control animals, 11 hM3Dq animals. All bars represent the mean of the data. Error bars are SEM. Images in **b** are representative of results observed across 5 (each WT, *Pdyn*^*Cre*^, *Dbh*^*Flp*^) and 11 (*Pdyn*^*Cre*^*;Dbh*^*Flp*^) animals. Scale bars are 100μm. Data point shape legend in **e** applies to data points in **e**-**g** and **j**-**k** and represent the genotypes of the mice. EZM - elevated zero maze; OFT - open field test; TH - tyrosine hydroxylase; LC - locus coeruleus; WT - wild-type; *Pdyn* - prodynorphin; *Dbh* - dopamine-β-hydroxylase.

## Discussion

### Unique features of ConVERGD and future directions

Current intersectional strategies are at risk of being outpaced by the need to target increasingly specific cell populations with increasingly complex payloads. To address this, ConVERGD is designed to facilitate the straightforward insertion of intact transgenes with minimal design constraints. This differs from current viral strategies utilized by the field, such as INTRSECT, which achieves intersectionality by splitting the transgene into separate exons placed within nested FLEx switches that are controlled by orthogonal recombinases^8,9^. Generating new INTRSECT tools requires complex genome engineering and codon optimization that must be customized for each new transgene^55^, a design strategy that may limit its widespread use and increase reliance on groups with specialized expertise in order to develop new versions. With ConVERGD, our intent is to expedite the development of diverse intersectional viral tools for a broader research community (Extended Data Fig. 2). Still, there are potential areas for improvement. For instance, while pairing ConVERGD with stronger promoters, such as CAG, increased intersectional gene expression, it also caused a modest increase in non-specific gene expression (Extended Data Fig. 1). We hypothesize that this may be caused by an inability of the ribozyme-guided cleavage to keep pace with mRNA production if transcription is too robust. While this issue can be circumvented by promoter choice, future iterations of ConVERGD may benefit from sophisticated efforts currently taking place in the field of synthetic biology to improve the function of ribozymes and other RNA switches^19,56,57^.

To our knowledge, ConVERGD is unique among neuroscience tools for its use of ribozymes to promote conditional gene expression, positioning it as a novel platform for future conditional tool design. While our work has focused on pairing ribozymes with recombinases to define complex intersectionality, ribozymes have also been shown to provide conditional expression using other strategies, such as morpholino binding^19^ or in combination with drugs like neomycin^58^ and tetracycline^59^. Thus, it should be feasible to expand ConVERGD’s design to include temporal intersection. Temporal control could also be achieved by combining ConVERGD tools with activity-dependent genetic labeling methods^60^, allowing end users to target distinct cellular populations based on their molecular identity and *in vivo* activity patterns. Ultimately, we reason that ConVERGD provides a solid starting point for building diverse intersectional vectors that encompass molecular and temporal dimensions, beyond what is possible with current strategies.

### Single-cell sequencing reveals molecular diversity in LC

While advances in single-cell sequencing technologies are rapidly improving our understanding of cellular diversity, these methods have not been uniformly applied to all brain regions. A prime example of this is the LC, a brain region traditionally considered to be molecularly homogeneous since almost all neurons within it produce NE. The fact that LC neurons collectively connect with most areas of the brain to release NE has further perpetuated a view that these cells uniformly and broadly modulate brain state. Yet, activation of LC neurons can promote diverse and even opposing behaviors related to arousal^31–33,61,62^, suggesting underlying heterogeneity within this circuit. While not explored in the context of the LC, other neuromodulatory cell types, such as dopaminergic and cholinergic neurons, can co-release neuromodulators and neurotransmitters that diversify their influence on downstream brain areas^63–68^. Expression of neuropeptides and neurotransmitters has been reported within the LC, both in classic histological studies using targeted probes, and more recently through bulk and single-cell transcriptomic studies performed on rodent LC neurons^24,26,69,70^. Still, we wondered if molecular heterogeneity within the LC could be further resolved using alternative approaches. For instance, we reasoned that collecting the entire LC cell body, rather than just nuclei, would provide a richer pool of mRNA for single-cell sequencing. Our initial efforts to collect LC neurons via standard dissociation and flow cytometry protocols resulted in extremely low LC cell viability (data not shown). Thus, we adopted electrophysiology-based techniques to prepare fresh brainstem tissue slices from which individual LC neurons were manually collected using pulled glass electrodes and suction. While laborious, this approach nicely complemented our choice to use Smartseq-2, which is also inherently low-throughput due to reagent cost and intensive protocol steps. From these experiments, we were able to profile 201 LC neurons (Fig. 5 and Extended Data Fig. 3), revealing differential expression of many neurotransmitters, neuropeptides, transporters, and receptors whose presence in the LC can now be more fully characterized using intersectional strategies such as ConVERGD. Additionally, we believe that the manual collection and sequencing strategy we report here could be useful for a variety of transcriptomic-based studies where cells of interest exist in low numbers. Ultimately, as experiments continue to uncover heterogeneity within neuronal populations like the LC, we will need easily adaptable tools to functionally characterize these cells. Development of ConVERGD allowed us to leverage insights gained from our LC single-cell sequencing experiments in order to perform this functional exploration.

## ACKNOWLEDGEMENTS

We thank Hollie Sanders and Kim Lowe for technical support, the St. Jude Vector Core Lab for generating ConVERGD AAVs, Geoff Neale and Scott Olsen in the St. Jude Hartwell Center for Biotechnology for guidance with sequencing, and Schwarz lab members for helpful feedback. We also thank Hongkui Zeng, Ali Cetin, Shenqin Yao, Thomas Zhou, and Marty T. Mortrud of the Allen Institute for sharing the N2c ^ΔG^-H2B-eGFP virus for transsynaptic tracing experiments and Guocai Zhong of Scripps Research, Florida for providing the initial sequence information for the T3H48 ribozyme. This work was supported by a NARSAD Young Investigator Grant from the Brain & Behavior Research Foundation (L.A.S), NIH grant 1DP2NS115764 (L.A.S), institutional funds from St. Jude Children’s Research Hospital (B.G.P, B.X, J.W.G, L.A.S.), and funding from the St. Jude Graduate School of Biomedical Sciences (A.C.H). Single-cell sequencing was performed at the Hartwell Center at St. Jude, which is supported in part by the National Cancer Institute of the National Institutes of Health under Award Number P30 CA021765.

## AUTHOR CONTRIBUTIONS

A.C.H. and L.A.S. conceived the project. A.C.H. designed ConVERGD, generated and tested viral constructs *in vitro* and *in vivo*, and performed rabies tracing and behavioral studies. B.G.P. assisted with the cloning of viral constructs and *in vitro* testing. B.X. performed sequencing analysis. J.W.G. piloted manual sequencing methods and collected cells for sequencing. P.C. and J.B.B. provided the protocol and starter virus for generating N2c-rabies. L.A.S. generated viruses, performed in situ hybridization experiments, *in vivo* testing and rabies tracing experiments, and supervised the project. A.C.H. and L.A.S. wrote and edited the manuscript with feedback from the other authors.

## Experimental Methods

### Key Resource Tables

| MOLECULAR CLONING |  |  |
| --- | --- | --- |
| Stable Competent E. coli | NEB | C3040H |
| Miniprep Classic kit | Zymo Research | D4054 |
| Gel DNA Recovery Kit | Zymo Research | D4008 |
| Nucleobond Extra Maxi EF kit | Machery-Nagel | 740424 |
| Sticky-end Ligase | NEB | M0370S |
| NEBuilder HiFi DNA Assembly Master Mix | NEB | E2621 |

| CELL CULTURE |  |  |
| --- | --- | --- |
| Neuro2a Cells | ATCC | CCL-131 |
| EMEM | GIBCO | 670086 |
| Pen/Step | GIBCO | 15140-122 |
| Fetal Bovine Serum | HyClone | SH30071.03 |
| Chamber slides | ThermoFisher | 154941PK |
| Lipofectamine 2000 | Invitrogen | 11668027 |

| LC SINGLE CELL SEQUENCING |  |  |
| --- | --- | --- |
| RNase inhibitor | Thermo Scientific | EO0384 |
| ERCC RNA spike-in | Invitrogen | 4456740 |
| Triton X-100 | Sigma Aldrich | 93443 |
| dNTPs | Invitrogen | 10297018 |
| 5X First Strand Buffer | Clontech | 639537 |
| DTT | Invitrogen | P2325 |
| Betaine | Sigma Aldrich | B0300-5VL |
| 1M MgCl<sub>2</sub> | Invitrogen | AM9530G |
| SmartScribe enzyme | Clontech | 639537 |
| 2X Kapa HiFi Hot Start Ready Mix | Kapa Biosystems | KK2601 |
| Lambda Exonuclease | NEB | M0262L |

| MOUSE LINES |  |  |
| --- | --- | --- |
| C57BL/6J (wildtype) | Jackson Laboratories | 000664 |
| *Dbh<sup>Flp</sup>* | Jackson Laboratories | 033952 |
| *Dbh<sup>Cre</sup>* | MMRRC | 032081-UCD |
| *Pdyn<sup>Cre</sup>* | Jackson Laboratories | 027958 |
| *Sun1-eGFP* | Jackson Laboratories | 021039 |
| *Ai14* | Jackson Laboratories | 007908 |

| SOFTWARE AND PROGRAMS |  |  |
| --- | --- | --- |
| SnapGene | GSL Biotech LLC | https://www.snapgene.com:443/ |
| FlowJo | Flowjo | https://www.flowjo.com |
| Bonsai | open ephys | open-ephys.org/bonsai |
| MATLAB | Mathworks | www.mathworks.com/products/matlab |
| R Studio | R | www.rstudio.com |
| Prism | Graphpad | www.graphpad.com/scientific-software/prism/ |
| FIJI | NIH | https://imagej.nih.gov/ij/ |
| TopScan | CleverSys | cleversysinc.com/CleverSysInc/csi_products/topscan-suite/#blank |

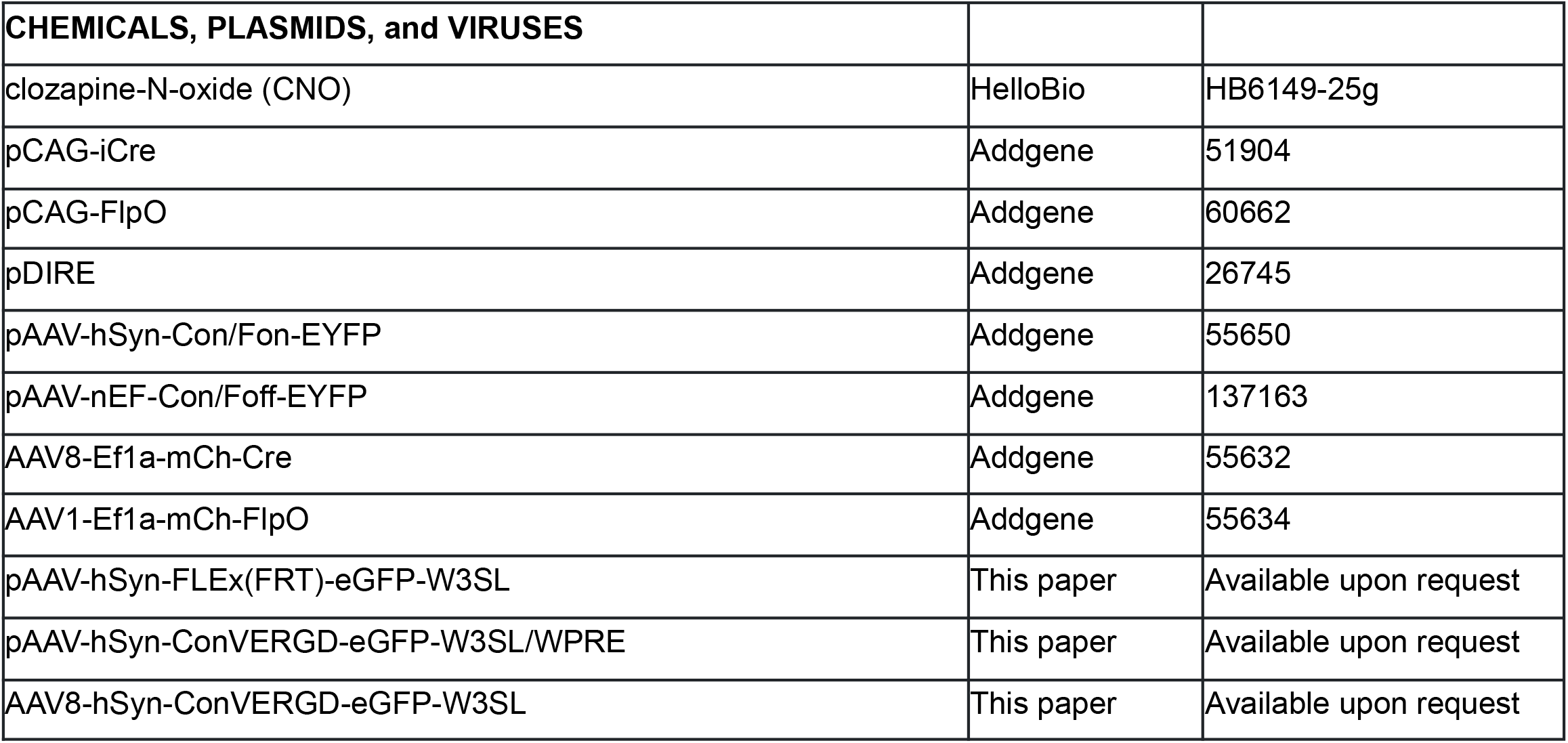
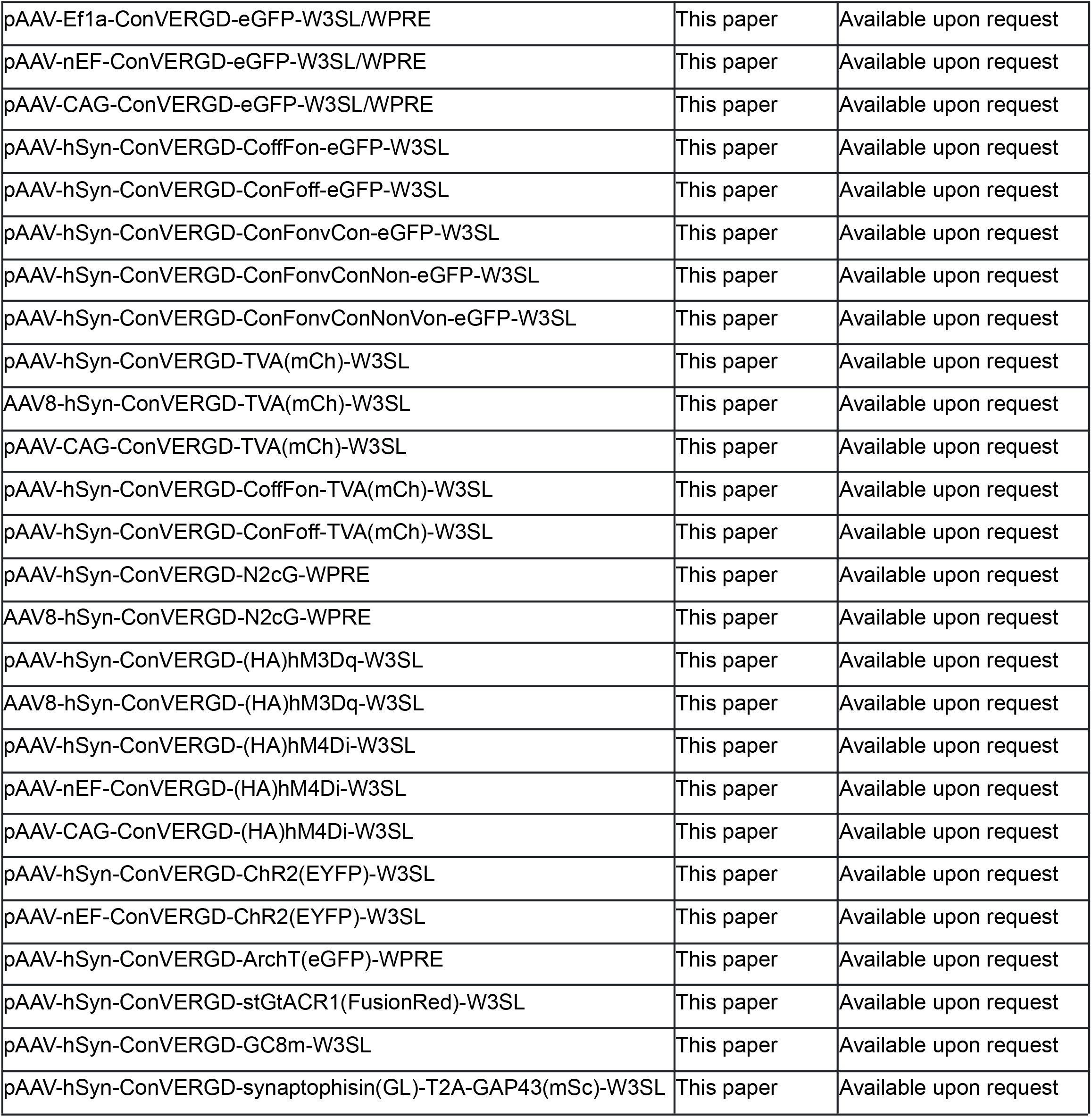

## Molecular Cloning

### Building ConVERGD

To begin building ConVERGD, we synthesized an AAV-backbone plasmid (pAAV-hSyn-FLEx(FRT)-eGFP-W3SL) (Azenta Life Sciences) containing the minimal human synapsin promoter (hSyn), an antisense eGFP coding region within a Flp-dependent FLEx-switch, the W3SL (WPRE3 (‘W’) followed by the SV40 late polyadenylation sequence with an upstream element (‘SL’)) 3’ posttranscriptional regulatory cassette, and robust multiple-cloning sites that provided easy component exchange. A custom oligo was synthesized (Azenta Life Sciences) that contained an hSyn promoter, a loxP-flanked (floxed) type III hammerhead ribozyme (T3H48)^19^, and an eGFP coding region within a Flp-dependent FLEx-switch. This ConVERGD-eGFP oligo was ligated into pAAV-hSyn-FLEx(FRT)-eGFP-W3SL backbone that had been previously digested with ApaI and HindIII restriction enzymes using Sticky-End ligation (NEB). This plasmid (called pAAV-hSyn-ConVERGD-eGFP-W3SL) was used for subsequent expansion of ConVERGD vectors.

### Testing ConVERGD backbone variants

To create ConVERGD with different promoters and 3’ posttranscriptional regulatory elements, the hSyn promoter was first excised from the pAAV-hSyn-ConVERGD-eGFP-W3SL backbone via NotI and KpnI restriction enzymes and replaced with the Ef1a promoter from Ef1a-SplitATG-eGFP-WPRE (gift from the Liqun Luo lab, Stanford University), the CAG promoter from pAAV-CAG-TVA(mChr)-WPRE (Addgene #48332, a gift from Liqun Luo) and the nEF promoter from a custom oligo designed based on the nEF promoter found in the map of nEF-Con/Fon-ChR2(ET/TC)-EYFP (Addgene #137139). The W3SL 3’ posttranscriptional regulatory element of all ConVERGD promoter variants was then excised with HindIII and BstEII restriction enzymes and replaced with the WPRE sequence from pAAV-CAG-TVA(mChr)-WPRE to create WPRE-containing ConVERGD variants.

### Expanding ConVERGD

To expand the ConVERGD toolkit to other Boolean expression logics, we synthesized custom oligos (Azenta Life Sciences) containing ConVERGD-eGFP (eGFP sequence and intersectional machinery) in CreOn;FlpOff, CreOff;FlpOn, and CreOn;FlpOn;vCreOn logics. pAAV-hSyn-ConVEGD-eGFP-W3SL was digested at NheI and HindIII to replace the original ConVERGD-CreOn;FlpOn logic with the additional Boolean logics listed above. To further expand ConVERGD to quadruple and quintuple intersectionality, we synthesized custom oligos Azenta Life Sciences) containing uniquely flanked ribozymes for CreOn;FlpOn;vCreOn;NigriOn and CreOn;FlpOn;vCreOn;NigriOn;VikaOn logics. ConVERGD-eGFP-ConFonvCon was then digested at EcoRI and HindIII sites to replace the original ConVERGD-ConFonvCon logic with the expanded logic cassettes.

### Exchanging ConVERGD payload

The eGFP coding region was excised from the pAAV-hSyn-ConVERGD-eGFP-W3SL backbone via digestion with EcoRI and SpeI restriction enzymes. Primers (IDT) designed via SnapGene were used to obtain a PCR product of the payload gene of interest with appropriate overhangs to be used in HiFi cloning. NEBuilder HiFi Assembly Master Mix was used with standard reagent ratios to insert the payload of interest into the linearized ConVERGD plasmid backbone. Transgenes inserted included: (HA)-hM3Dq (from Addgene #125147, a gift from Christopher Ritchie), (HA)-hM4Di (from Addgene #125146, a gift from Christopher Ritchie), ChR2(YFP) (from Addgene #20298, a gift from Karl Deisseroth), ArchT-eGFP (from Addgene #28307, a gift from Edward Boyden), stGtACR1(FusionRed) (from Addgene #105678, a gift from Ofer Yizhar), TVA-(mCherry) (from Addgene #48332, a gift from Liqun Luo), N2cG (from Addgene #73481, a gift from Thomas Jessell), GCaMP8m (from Addgene #162378, a gift from the GENIE Project), Synaptophysin-GreenLantern-GAP43-mScarlet (generated in the Schwarz lab).

### Comparing the genetic size of ConVERGD and INTRSECT intersectional machinery

To directly compare the genetic space needed for intersectional machinery between ConVERGD and INTRSECT constructs, we added the basepairs used by the intersectional components of each strategy. As backbones of each strategy differed, we did not include promoters, posttransciptional regulatory elements, restriction sites, or spacer regions not directly related to FLEx switches or intronic regions. ConVERGD numbers included only recombinase sites, spacer sequences between recombinase sites (if present), and ribozymes. INTRSECT numbers were based on double intron intersectional constructs and included only recombinase sites, spacer sequences between recombinase sites, intronic sequences (acceptors, donors, and spacers). Numbers were derived from sequences available at https://web.stanford.edu/group/dlab/optogenetics/sequence_info.html#intrsect.

### Virus production

AAVs were synthesized by the St. Jude Viral Vector Core using standard protocols or purchased directly from Addgene. RABV-CVS-N2c^ΔG^-H2B-eGFP rabies virus was propagated in the Schwarz lab from an aliquot obtained from the Allen Institute for Brain Science using cell lines and protocols described in Reardon et al.^47^

### *In vitro* testing

Mouse neuroblastoma (N2a) cells (ATCC) were co-transfected with ConVERGD and recombinase-expressing plasmids (Cre, Flp, and Cre+Flp (pDIRE)). Cells were cultured in a 10cm dish (FisherBrand) with EMEM (GIBCO) + 5% pen/strep (GIBCO) + 10% fetal bovine serum (FBS) (HyClone). Upon reaching 80-90% confluency, cells were split, passaged, and used to seed either CC2-treated 8-chamber slides (Thermo Fisher) for qualitative analyses or cell-culture treated 24-well plates (Costar) for collection and quantitative FACS analyses. 4500-8000 cells were used to seed each well of the slides and the 24-well plates. One day after plating, cells were transfected using Lipo2000 (Invitrogen) with standard reagent ratios. For chamber slide transfections, 50ng of plasmid DNA were used (e.g. 50ng ConVERGD vector with 50ng recombinase vector). For 24-well plate transfections, 250ng of plasmid DNA were used for all samples (e.g. 250ng ConVERGD vector with 250ng recombinase vector). Two days after transfection, cells in the chamber slides were fixed with 4% paraformaldehyde (PFA) in PBS and stained with the appropriate antibody, if necessary. All fixed slides were counterstained with 1:10,000 of 5 mg/ml 2 1 4′,6-diamidino-2-phenylindole (DAPI; Sigma-Aldrich) in PBS. Fixed slides were imaged with a Leica DM6 automated fluorescence microscope for qualitative analysis. For quantitative FACS analysis, cells in 24-well plates were trypsinized three days post-transfection and collected into individual 1.5ml tubes. Cells were pelleted at 200g for 5min, resuspended in 200ul of 1x PBS, and transferred to filtered FACS tubes (Fischer) on ice. DAPI was added as a counterstain to exclude dead cells immediately prior to FACS analysis (BD FACSAria Fusion). FlowJo was used to analyze and quantify the results. Cells were positively gated for high-density populations plotted by side scatter (SSC-A) and forward scatter (FCS-A) (to isolate cells from debris), positively gated for DAPI negative cells (to isolate live cells from dead cells), positively gated for high-density populations plotted by FSC-A and SSC-W (to isolate single cells), and finally, positively gated for GFP^+^ cells (to isolate GFP^+^ cells). Samples with less than 7,500 live, single cells were excluded from analysis. Normalized control values were calculated by averaging the median fluorescent intensity (MFI) readings of all control samples for each constructs. Fold induction values reported represent fold change of each sample from these normalized control values. All quantified transfection experiments were performed at least 3 times.

### Animals

Adult (7-37 weeks) male and female mice, C57BL/6J wild-type (JAX# 000664), *Dbh*^*Flp*^ (JAX# 033952), *Dbh*^*Cre*^ (MMRRC# 032081-UCD), *Pdyn*^*Cr*e^ (JAX# 027958), *Sun1-eGFP* (JAX# 021039), and *Ai14* (JAX# 007908) were housed up to 5 mice per cage and kept on a 12-hour light/dark cycle (lights on at 06:00) with *ad libitum* food and water. Experimental protocols were approved by and conformed to St. Jude Children’s Research Hospital IACUC and conform to US National Institutes of Health guidelines.

### Stereotaxic Surgery

Mice were anesthetized and maintained on isoflurane and prepared for surgery following institutional guidelines. Craniotomies were performed to deliver virus directly to targeted brain regions via a microinjection pump (World Precision Instruments) and pulled glass electrode. The locus coeruleus was targeted with the following coordinates from lambda: -0.8mm (posterior), +/-0.8 (lateral), 3.2mm (ventral from the surface of the brain). ConVERGD viruses were diluted to reduce viral leak expression. We found a titer of ∼8E+11gc/mL worked best for AAV8-hSyn-ConVERGD-eGFP-W3SL and AAV8-hSyn-ConVERGD-(HA)hM3Dq-W3SL. For ConVERGD-eGFP + recombinase virus experiments, a working titer of ∼7.87E+11gc/mL was used for ConVERGD-eGFP for all injections. For expression comparisons between ConVERGD and INTRSECT, a working titer of ∼7.87E+11gc/mL was used for all injections of both viruses. For rabies tracing experiments, AAV8-hSyn-ConVERGD-TVA(mChr)-W3SL and AAV8-hSyn-ConVERGD-H2B-P2A-N2cG-WPRE were mixed for a working titer of 3.78E12 gc/mL for TVA and 3.81E12 gc/mL for N2c. The titer of RABV-CVS-N2c^ΔG^-H2B-eGFP was ∼2.1 × 10^8 infectious particles per ml based on serial dilutions of the virus stock followed by infection of the 293-TVA800 cell line.

### Behavioral Assays

Adult male and female mice (10F, 16M) 10-19 weeks old at the time of behavior were used for these experiments. All mice were bilaterally injected with 350nl of AAV8-hSyn-ConVERGD-(HA)hM3Dq-W3SL into the LC. Mice were divided into control and experimental (hM3Dq-expressing) groups. The control group consisted of WT (2F, 3M), *Pdyn*^*Cre*^ (2F, 3M), and *Dbh*^*Flp*^ (2F, 3M) mice. The experimental (hM3Dq-expressing) group consisted of *Pdyn*^*Cre*^*;Dbh*^*Flp*^ (4F, 7M) mice. Histology was performed on all animals after the completion of behavioral studies to confirm the presence (for experimental mice) or absence (for control mice) of hM3Dq in the LC. Behavioral data for control genotypes (WT, *Pdyn*^*Cre*^, and *Dbh*^*Flp*^ mice) was combined after ordinary one-way ANOVA tests with a post-hoc Tukey’s multiple comparisons test showed no statistical difference between genotypes. EZM control group p-values: % of time in open region - WT vs *Pdyn*^*Cre*^ vs *Dbh*^*Flp*^: 0.3656 (ANOVA), WT vs *Pdyn*^*Cre*^: 0.6926 (adjusted, Tukey’s), WT vs *Dbh*^*Flp*^: 0.3358, *Pdyn*^*Cre*^ vs *Dbh*^*Flp*^: 0.7975 (adjusted, Tukey’s); % of distance in open region - WT vs *Pdyn*^*Cre*^ vs *Dbh*^*Flp*^: 0.9578 (ANOVA), WT vs *Pdyn*^*Cre*^: 0.9599 (adjusted, Tukey’s), WT vs *Dbh*^*Flp*^: 0.9710 (adjusted, Tukey’s), *Pdyn*^*Cre*^ vs *Dbh*^*Flp*^: 0.9991 (adjusted, Tukey’s). OFT control group p-values: % of time in center - WT vs *Pdyn*^*Cre*^ vs *Dbh*^*Flp*^: 0.3320 (ANOVA), WT vs *Pdyn*^*Cre*^: 0.8471 (adjusted, Tukey’s), WT vs *Dbh*^*Flp*^: 0.7131 (adjusted, Tukey’s), *Pdyn*^*Cre*^ vs *Dbh*^*Flp*^: 0.9682 (adjusted, Tukey’s); % of time in corners - WT vs *Pdyn*^*Cre*^ vs *Dbh*^*Flp*^: 0.4785 (ANOVA), WT vs *Pdyn*^*Cre*^: 0.8764 (adjusted, Tukey’s), WT vs *Dbh*^*Flp*^: 0.4517 (adjusted, Tukey’s), *Pdyn*^*Cre*^ vs *Dbh*^*Flp*^: 0.7385 (adjusted, Tukey’s).

### Elevated zero maze (EZM)

The EZM apparatus was a 7cm wide, circular track 52cm in diameter elevated 60cm off the ground. The track contained two closed regions on opposite sides of the maze with 14cm tall opaque walls that each ran ¼ the circumference of the track. At least 3 weeks after bilateral injection of AAV8-hSyn-ConVERGD-(HA)hM3Dq-W3SL into the LC, mice were handled for 4 days for acclimation. Day 1 of handling consisted of the experimenter placing their hands in the home cage for 5 minutes. On day 2, the experimenter held each mouse for 5 minutes. On day 3, mice were weighed, scruffed, and given an intraperitoneal (IP) saline injection of the same volume that would be injected of the CNO solution on the test day. On day 4, mice were weighed, scruffed, and given an IP injection of saline 30 minutes prior to being placed on the EZM for ∼12 minutes. On the test day, mice were acclimated to the testing room for at least 15 minutes prior to being weighed, scruffed, and given an IP injection of 5mg/kg clozapine-N-oxide (CNO; HelloBio). 30 minutes after CNO injection, mice were placed on the EZM at the border of the open and closed region facing the open region. Mice were video recorded from above for the entirety of the 10min trial. Centroid tracing data was acquired with a custom Bonsai program and analyzed using a custom MATLAB script. To determine the time spent in each region, Bonsai was used to draw ROIs around each open region, and the amount of time the ROI was triggered was recorded in real time. This method was very sensitive and counted any triggering of the ROI, including head pokes into the open region, as time spent in the open region. However, the threshold on the video analysis was set such that the tail was not visible by the ROI. This ensured exclusion of false triggers that could be caused by the mouse’s tail lying across the closed/open threshold. To calculate the distance traveled in each region, MATLAB was used to analyze centroid data obtained from the Bonsai tracking. ROIs were drawn around the closed regions and the cumulative distance between centroid points within these regions was calculated.

### Open field testing (OFT)

The open field chamber consisted of a 16inx16in plastic chamber with 14in tall transparent walls enclosing the chamber. Mice were video recorded from above for the entirety of the 10min trial. The same mice used in the EZM experiments were used for OFT. One week after EZM testing with CNO, mice were acclimated to the OFT room for at least 15 minutes, weighed, scruffed, and given an IP injection of 5mg/kg CNO 30 minutes prior to being placed in the open field chambers. CleverSys TopScan software was used to track and analyze OFT behavior.

## Single-cell RNA sequencing

Adult *Dbh*^*Cre*^*;Ai14* mice (∼8-20 weeks old) were euthanized with tribromoethanol (Avertin). Fresh brain tissue was immediately extracted and placed in ice-cold aCSF (2.5 mM KCl, 7 mM MgCl_2_, 0.5 mM CaCl_2_, 1.3 mM NaH_2_PO_4_, 110 mM choline chloride, 25 mM NaHCO_3_, 1.3 mM sodium ascorbate, 20 mM glucose, 0.6 mM sodium pyruvate, bubbled in 95% O2 /5%CO2). Submerged brain tissue was placed on a VT1200 vibratome (Leica) and 200 μm coronal slices that included the LC were collected. Slices were transferred to a 32°C holding chamber containing oxygenated aCSF, where they recovered for at least 1 hour. Cells were visualized using an Olympus BX-51 fluorescence microscope, and tdTomato-expressing LC neurons were manually extracted from the slice using glass pipettes pulled from borosilicate glass (World Precision Instruments) to create an ∼50 μm tip. The ends of glass pipette tips containing picked cells were snapped into individual wells of a 96 well plate containing 4μl lysis buffer: 0.1μl RNase inhibitor (Thermo Scientific), 0.1μl ERCC RNA spike-in (Invitrogen), 0.02μl 10% Triton (Sigma), 1μl 10mM dNTP (Invitrogen), 0.1μl 100μM dT (IDT; 5’-AAGCAGTGGTATCAACGCAGAGTACTTTTTTTTTTTTTTTTTTTTTTTTTTTTTTVN), and 2.68μl H_2_O. Following collection, plates were spun down, sealed, and stored at -80°c. Samples next underwent primer annealing (72°c for 3min) before placing on ice. Next, 6μl reverse transcription reagent was added to each sample: 0.14μl H_2_O, 0.25μl RNase inhibitor, 2μl 5X First Strand Buffer (Clontech), 0.5μl 100mM DTT (Invitrogen), 2μl 5M Betaine (Sigma), 0.06μl 1M MgCl2 (Invitrogen), 0.1μl 100μM TSO (Qiagen; 5’-AAGCAGTGGTATCAACGCAGAGTACATrGrG+G), 0.95μl SmartScribe enzyme (Clontech). Samples were incubated at 42°c for 90min, 70°c for 5min, then held at 4°c. DNA preamplification solution (15μl) was next added to each sample: 2.1375μl H_2_O, 12.5μl 2X Kapa HiFi Hot Start Ready Mix (Kapa Biosystems), 0.25μl 10μM IS_PCR Primer (IDT; 5’-AAGCAGTGGTATCAACGCAGAGT), 0.1125μl Lambda Exonuclease (NEB). Samples were incubated at 37°c for 30min, 95°c for 3min, 21 cycles of [98°c for 20s, 67°c for 15sec, 72°c for 4min], 72°c for 5min, and then held at 4°c. DNA cleanup, quantification, dilution, tagmentation, and barcoding were performed by the Hartwell Center at St. Jude Children’s Research Hospital using standard methods. Libraries were sequenced on a HiSeq4000 or MiSeq Sequencing Systems (Illumina) using 2 X100-bp or 2 × 75-bp ends, respectively. Sequences were de-multiplexed using bcl2fastq and aligned to mouse mm10 genome (with Cre and tdTomato genes added). Paired-end reads from Illumina were trimmed for adapters by cutadapt(v1.15) and aligned to mm9 (MGSCv37, NCBI) by STAR(v2.5.2b). LiftOver was performed on GENCODE(vM9) gene database (from mm10 to mm9) by CrossMap(v0.1.5) to improve gene annotation. Aligned reads were then counted by HTSEQ (v 0.6.1p1) for genes annotated in GENCODE database. Trimmed reads were also run through FastQ Screen(v0.9.5) against Human, Mouse, Adapters, Univec databases (Dec 2016) to check for contamination and doublets. Samples with larger unique hits to human over mouse genomes were considered putative doublets and excluded from analysis. We further excluded samples with less than 1M reads and those that did not express *Dbh*. We then conducted an unbiased analysis to identify the top 100 genes that appeared most frequently across LC samples. Next, we downloaded gene lists from Gene Ontology (GO) for selected terms and plotted heatmaps (Z score) for the top expressed genes in each GO term using the ComplexHeatmap R package.

## Histology

### In situ hybridization

Mice were euthanized with tribromoethanol (Avertin) and fresh brain tissue was harvested and immediately mounted in OCT for cryosectioning. 16-20 μm tissue sections containing the LC were collected for RNAscope-based fluorescence in situ hybridization (ACDBio). Samples were labeled using the RNAscope Multiplex Fluorescent Reagent Kit v2 (ACDBio), and following the supplied protocol. The following probes were used: Calca-C1 (420361) Dbh-O1-C3 (464621-C3), Gad2-C1 (439371) Pdyn-C1 (318771), Penk-C1 (318761), Slc17a6-C2 (319171-C2), Slc17a7-C2 (416631). Probe signal was developed using Opal dyes (Opal 520 and 570 Reagent Pack, Perkin Elmer) at a dilution of 1:1000 and counterstained with DAPI. Fluorescent images were taken using a Leica LSM780 laser scanning confocal microscope.

### Combined in situ hybridization and immunostaining

Mice were euthanized with tribromoethanol (Avertin) and fresh brain tissue was harvested and immediately mounted in OCT for cryosectioning. 20 μm tissue sections containing the LC were collected for Molecular Instruments-based hybridization chain reaction (HCR) *in situ* hybridization, Samples were labeled using custom probes (*Dbh-B2* and *Pdyn-B5*), and following the supplied protocol. Following RNA amplification and labeling, slides were washed with PBS (3 × 15min), followed by blocking solution (PBS solution containing 0.05% Triton X-100 and 3% BSA) for 1 hour at room temperature. Slides were then incubated overnight at room temperature with GFP-rb antibody (1:500). The following day, slides were washed with a PBS solution containing 0.1% Tween 20 before application of secondary antibodies (1:500) for 2 hours at room temperature. All sections were counterstained in DAPI prior to mounting (Fluoromount-G; SouthernBiotech). Fluorescent images were taken using a Leica LSM780 laser scanning confocal microscope.

### Immunostaining

Mice were euthanized with tribromoethanol (Avertin) and transcardially perfused with PBS and 4% PFA. Brain tissue was removed, placed in 4% PFA at 4°C overnight, and then equilibrated to a 30% sucrose solution at 4°C until they settled at the bottom of the container (2-4 overnights). Brains were cryo-sectioned (Leica) at 50μm and collected on Superfrost+ slides, Superfrost slides, or directly into wells containing 1x PBS. Sections collected on Superfrost slides were then floated with 1x PBS and transferred to wells containing 1x PBS. All sections were washed 3 times in 1x PBS for 10 minutes. Slide attached sections were stained in the appropriate primary antibody at 4°C for 3 overnights. Floating sections were stained with the appropriate primary antibody at room temperature overnight with shaking. All primary antibodies were used at a 1:1000 dilution in a PBS solution containing 2% NDS and 0.2% Triton X-100. All sections were stained with the appropriate secondary antibody at room temperature with shaking for 2 hours. Secondary antibodies were diluted 1:500 in a PBS solution containing 2% NDS. All sections were counterstained in DAPI prior to mounting (Fluoromount-G; SouthernBiotech). ConVERGD-eGFP, INTRSECT-YFP, and *Pdyn*^*Cre*^; *Sun1-eGFP* samples were stained with GFP-Ch and Th-Rb. For cFos staining, every other section of ConVERD-(HA)hM3Dq injected brains were stained with TH-Rb and HA-Ms or TH-Ms and cFos-Rb. Because of the overlap of antibodies against TH, we did not stain against TH, HA, and cFos together. Fluorescent images were taken using a Nikon C2 or Leica LSM780 laser scanning confocal microscope or a Leica DM6 automated fluorescence microscope.

| Antibody | Species | Dilution | Source |
| --- | --- | --- | --- |
| TH | Rabbit | 1:1000 | EMD Millipore (ab152) |
| TH | Mouse | 1:1000 | EMD Millipore (MAB318) |
| HA.11 | Mouse | 1:1000 | BioLegend (901513) |
| cFos | Rabbit | 1:1000 | Synaptic Systems (226008) |
| GFP | Chicken | 1:1000 | Aves Labs (GFP3717982) |
| GFP | Rabbit | 1:1000 | Invitrogen (A11122) |
| RFP | Rabbit | 1:1000 | Rockland (600-401-379) |
| Alexa Fluor 647<br>anti-rabbit IgG | Donkey | 1:500 | JacksonImmuno (711-605-152) |
| Alexa Fluor 488<br>anti-rabbit IgG | Donkey | 1:500 | JacksonImmuno (711-545-152) |
| Alexa Fluor Cy3<br>anti-rabbit IgG | Donkey | 1:500 | JacksonImmuno (711-165-152) |
| Alexa Fluor 647<br>anti-mouse IgG | Donkey | 1:500 | JacksonImmuno (711-605-150) |
| Alexa Fluor 488<br>anti-mouse IgG | Donkey | 1:500 | Invitrogen (A32766) |
| Alexa Fluor 488<br>anti-chicken IgG | Donkey | 1:500 | JacksonImmuno (711-545-155) |

### Fluorescent image analysis

To compare fluorescently labeled cells between ConVERGD-eGFP and INTRSECT-EYFP, 6 50μm sections spanning the anterior-posterior axis of the LC were selected from at least 3 mice per genotype (*Pdyn*^*Cre*^, *Dbh*^*Flp*^, *Pdyn*^*Cre*^, *Dbh*^*Flp*^) per virus injection group. All sections were immunostained against GFP as described above, and images were taken with a Leica LSM780 laser scanning confocal microscope. Fluorescently labeled cells within these images were manually counted. To quantify the rabies-tracing experiments, brains were cryosectioned at 50μm, and every other section throughout the entire brain was collected onto Superfrost+ slides. Images of every other collected section were taken using a 5X objective on a Leica DM6 fluorescent microscope. For each brain, GFP^+^ nuclei were manually quantified across every image, totaled for all images, and then each image was assigned a percentage of the total counted GFP^+^ nuclei within that brain. Fluorescence images were then manually registered to the corresponding image in a reference atlas (Paxinos and Franklins). Using Rstudio, data were grouped by the registered atlas image number, and an average percentage of the total number of labeled cells was determined for each registered image section. Peaks in the average labeled cell count of at least 3% of total counted cells were chosen to obtain example images from the major input regions. Example input images were taken with a Nikon C2 fluorescent confocal microscope. To visualize axon projections, we analyzed sections that had previously been collected for rabies quantification. These sections were immunolabeled with an anti-RFP antibody to amplify the signal from the expression of AAV8-hSyn-ConVERGD-TVA(mCh)-W3SL. Qualitative images of axon projections within regions of interest were taken using a Nikon N2c fluorescent confocal microscope. Axon projection patterns to brain regions represented in Fig 7 were observed in multiple samples.

## Data Availability

RNA sequencing data is deposited in the NCBI Gene Expression Omnibus (GEO) database with accession code GSE224285. Other data from the paper (behavior, FACS, histology images) is available from the corresponding author upon reasonable request.

---

**Extended Data Fig. 1.**
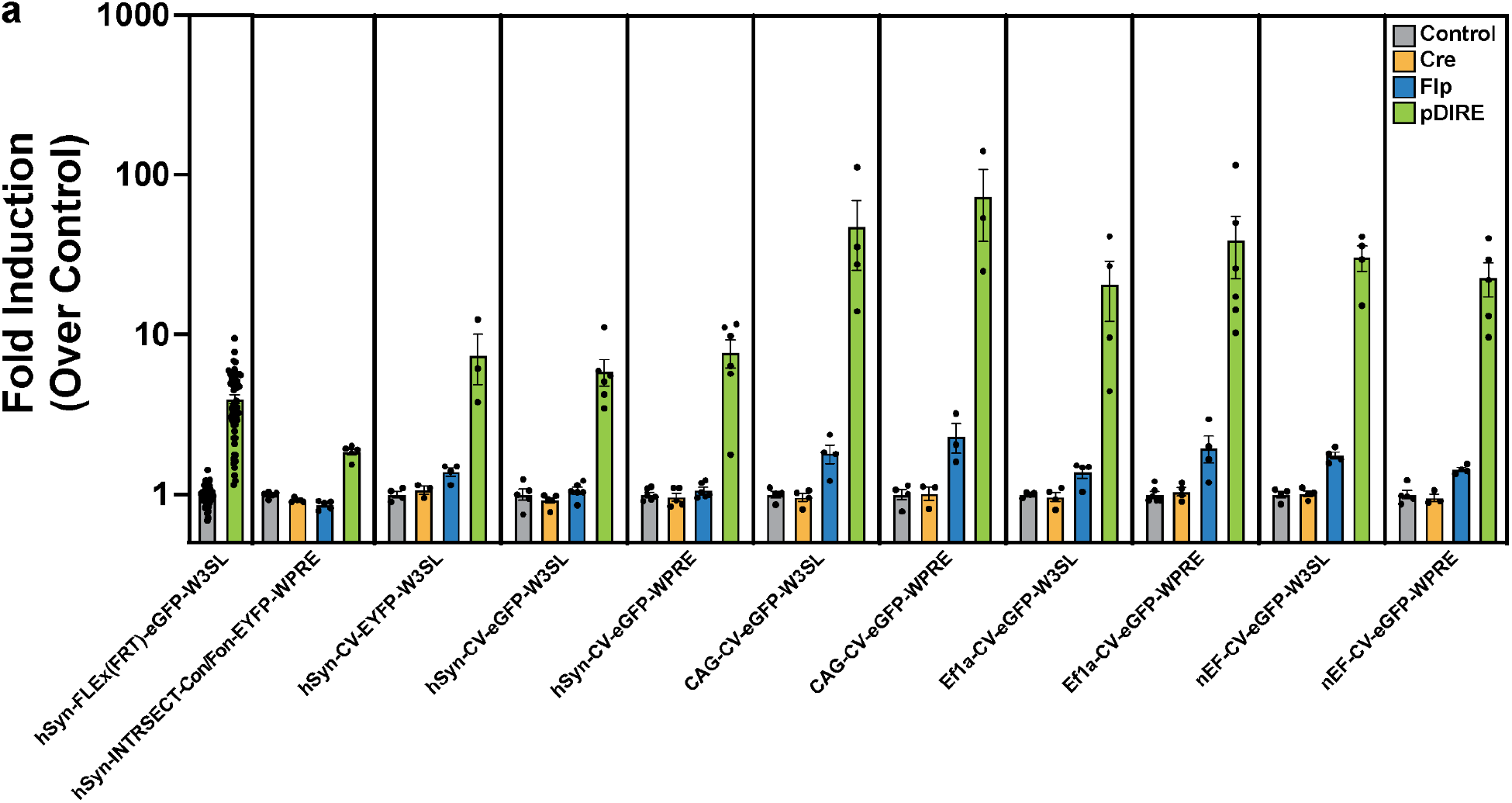
Comparison of ConVERGD-based constructs across conditions. **a**, FACS quantification of N2a cells co-transfected with an EYFP- or eGFP-expressing plasmid alone (Control, grey bars) or with recombinase plasmids expressing Cre (yellow bars), Flp (blue bars), or Cre and Flp (pDIRE, green bars). ConVERGD was tested in pAAV backbones containing different promoters and 3’ posttranscriptional regulatory elements. Results represent data from at least 3 separate transfection experiments. Data points represent fold induction values of individual transfections compared to the average control median fluorescent intensity (MFI) for each construct. Bars represent the mean of all experiments. Error bars are SEM. CV - ConVERGD.

**Extended Data Fig 2.**
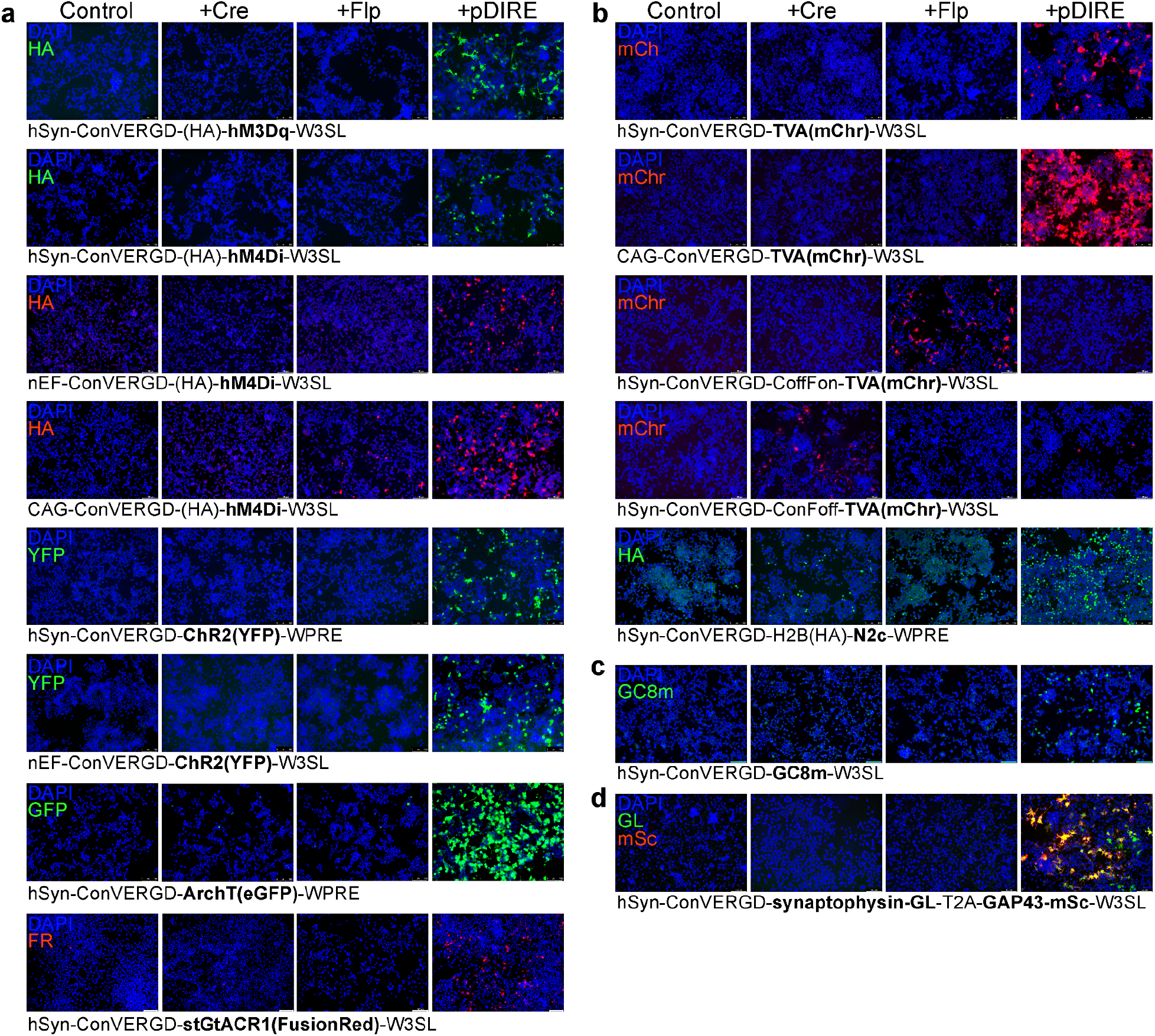
ConVERGD-based constructs are easily amenable and allow specific expression of diverse transgenes. **a**, ConVERGD-based toolkit for modulating neuronal activity. **b**, ConVERGD-based toolkit for trans-synaptic rabies tracing. **c**, ConVERGD-based construct for *in vivo* calcium imaging (GCaMP8m; GC8m). **d**, ConVERGD-based construct for a dual-expressing transgene that labels pre-synaptic sites and axons (synaptophysin-GreenLantern and GAP43-mScarlet). All images show transfected N2a cells counterstained with DAPI (blue) and are representative of results observed across at least 2 separate transfections. FR - FusionRed; mChr - mCherry; GL - GreenLantern; mSc - mScarlet.

**Extended Data Fig. 3.**
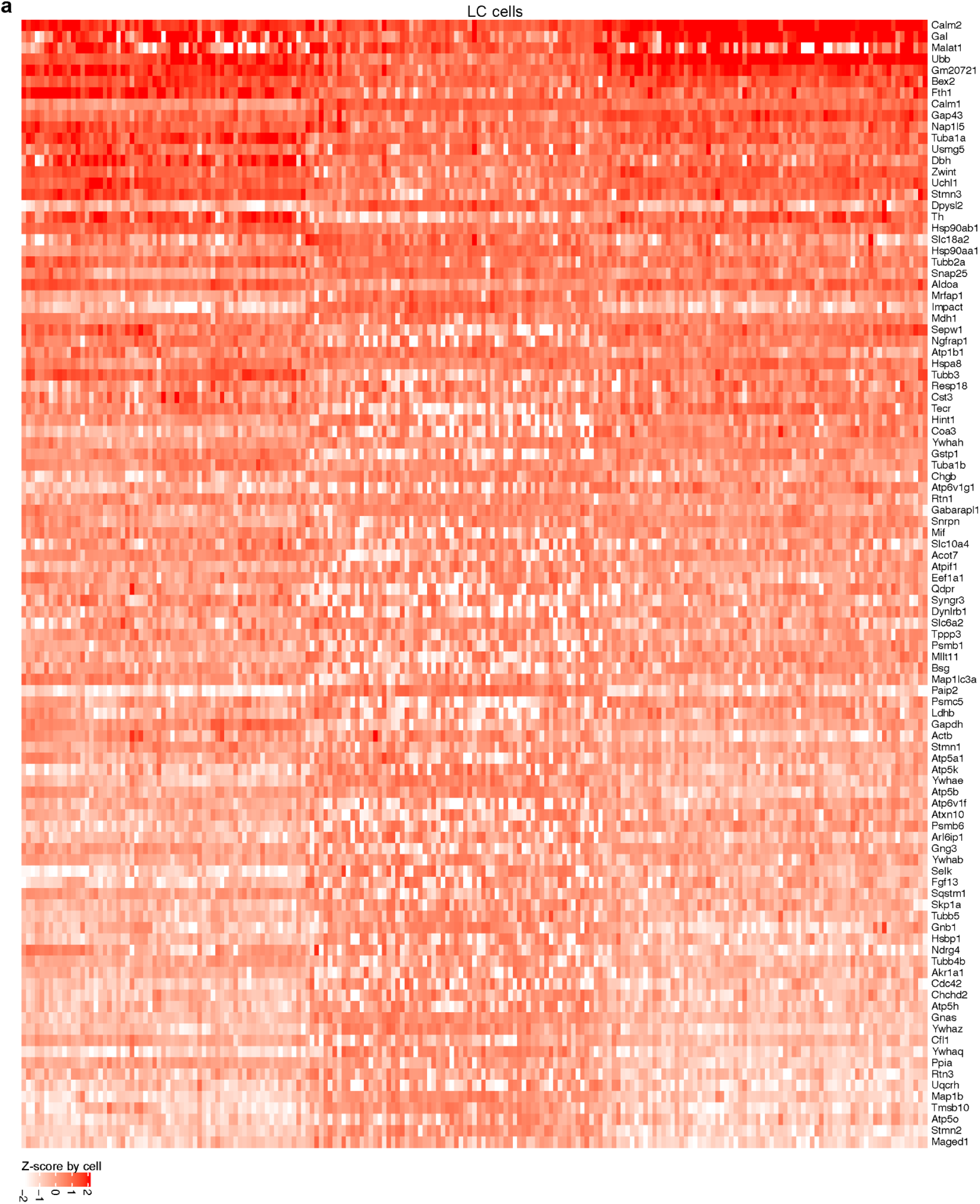
Top 100 most frequently detected genes in LC neurons using a Smart-seq2-based sequencing platform. **a**, Heatmap of scaled (by cell) transcript abundance (transcripts per million, TPM) for the top 100 genes most frequently detected in 201 LC neurons by single-cell transcriptomic sequencing.

**Extended Data Fig. 4.**
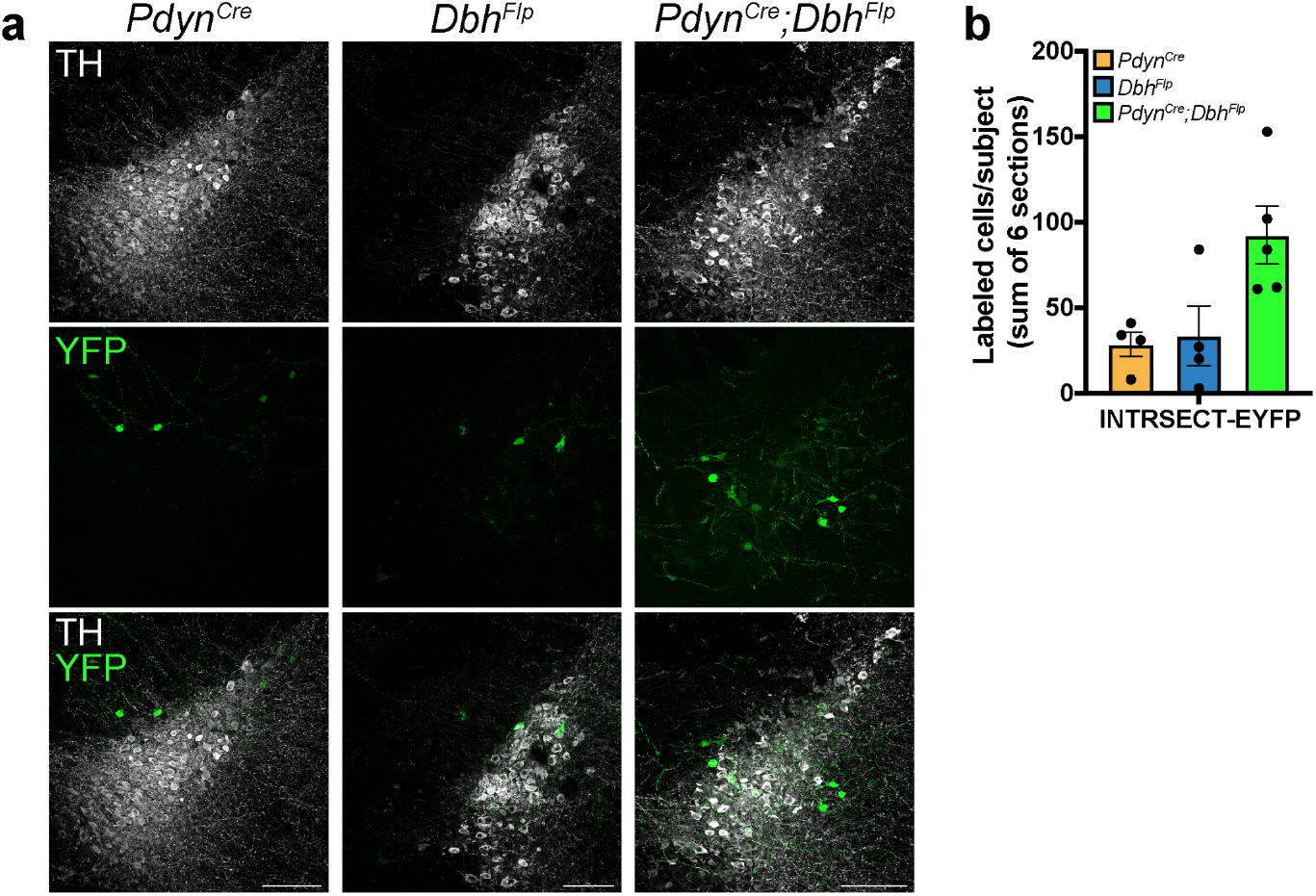
Increased single-recombinase induced expression observed with INTRSECT. **a**, Representative images showing the LC (TH, white) of mice injected with AAV-hSyn-INTRSECT-Con/Fon-EYFP. All genotypes showed some level of YFP (green) expression. **b**, Quantification of INTRSECT-EYFP labeled cells in and around (∼200μm radius) the LC across different genotypes. n = 4 (*Pdyn*^*Cre*^), 4 (*Dbh*^*Flp*^), and 5 (*Pdyn*^*Cre*^*;Dbh*^*Flp*^). Points represent the averaged cell counts across 6 50μm LC brain sections. Bars represent the mean of the data. Error bars are SEM. All sections were immunostained against GFP. Images in **a** are representative of results observed across 4 (*Pdyn*^*Cre*^), 4 (*Dbh*^*Flp*^), and 5 (*Pdyn*^*Cre*^*;Dbh*^*Flp*^) animals. Scale bars are 100μm. TH - tyrosine hydroxylase; LC - locus coeruleus; *Pdyn* - prodynorphin; *Dbh* - dopamine-β-hydroxylase.

**Extended Data Table 1.** Genetic space needed for intersectional machinery. Estimate of base pairs occupied by intersectional genetic machinery for ConVERGD and double-intronic INTRSECT strategies. Totals include recombinase sites, spacer regions between FLEx-oriented recombinase sites, intronic acceptor and donor regions, and ribozymes, where applicable. Since the vector backbone of each strategy differed, ITRs, promoters, restriction digest sites, transgene coding regions, and 3’ UTRs were not included in these counts. 3x - triple intersectional (CreANDFlpANDvCre); 4x - quadruple intersectional (CreANDFlpANDvCreANDNigri); 5x - quintuple intersectional (CreANDFlpANDvCreANDNigriANDVika); “---” - represents a lack of 4x and 5x INTRSECT constructs.

|  | **Intersectional Strategy** |  |
| --- | --- | --- |
| **Expression Logic** | **ConVERGD (bp)** | **INTRSECT (bp)** |
| CreOn/FlpOn | 448 | 666 |
| CreOff/FlpOn | 564 | 666 |
| 3x-AND | 508 | 901 |
| 4x-AND | 640 | --- |
| 5x-AND | 800 | --- |

